# Modulating Neurotoxic Effects of Prenatal Chlorpyrifos Exposure Through Probiotic and Vitamin D Gestational Supplementation: Unexpected Effects on Neurodevelopment and Sociability

**DOI:** 10.1101/2024.11.04.621795

**Authors:** Mario Coca, Cristian Perez-Fernandez, Ana C. Abreu, Ana M. Salmerón, Miguel Morales-Navas, Diego Ruiz-Sobremazas, Teresa Colomina, Ignacio Fernández, Fernando Sanchez-Santed

## Abstract

Autism is a neurodevelopmental disorder characterized by impairments in sociability and communication. Prenatal exposure to Chlorpyrifos has been associated with autism-like behaviors in preclinical models. Interest has grown in the gut-brain axis and the role of microbiota modulation through dietetic supplementation to reduce this ASD-like phenotype. This study examines the effects of prenatal CPF exposure in Wistar Rats and assesses the potential of gestational probiotic and vitamin D supplementation to mitigate these effects in offspring. CPF exposure significantly impaired sociability in adolescence, and supplementation did not reverse these deficits. However, in control animals, supplementation induced neurodevelopmental changes, including alterations in metabolic status, the pattern of expression of ASD-related genes, the regulation of oxytocin and vasopressin receptors, and the GABAergic system in the brain. Additionally, supplementation accelerated overall development, increased ultrasonic vocalization emission and modified the typical responses to social novelty. CPF exposure blocked most of these effects at both behavioral and molecular levels. While supplementation did not block CPF-induced impairments, CPF exposure altered the observed effects of supplementation in controls, possibly indicating shared molecular mechanisms. These findings highlight the need for further research into the safety of probiotic and vitamin D supplementation during pregnancy.

## 1. INTRODUCTION

Autism spectrum disorder (ASD) is a complex developmental disorder characterized by varying degrees of impairment in sociability, verbal and non-verbal communication, and the presence of repetitive and stereotyped behavioral patterns, activities, or interests (Lord et al., 2018). Additionally, it involves alterations in other areas, such as motor skills, attention, or inhibitory control (Schmitt et al., 2018; Biosca-Brull et al., 2021). These symptoms vary depending on factors such as age, sex, and developmental level, with males being more prone to autism (García-Franco et al., 2019). Although the exact cause of ASD remains unknown, some researchers have suggested that abnormal GABA maturation may play a significant role in its pathology (Zhao et al. 2021). During postnatal development, the GABAergic system switches from excitatory to inhibitory signaling, driven by the upregulation of KCC2 expression and the downregulation of NKCC1 (Peerboom and Wierenga, 2021). Disruptions in this developmental milestone, which affect the balance between excitatory and inhibitory (E/I) balances, have been observed in autism (Eissa et al., 2018; Pozzi et al., 2020). ASD has also been linked to alterations in several other neurotransmitter and neuromodulator systems, including oxytocinergic, vasopressinergic, glutamatergic, and serotonergic systems. Additionally, several other molecular pathways — including those involving BDNF and PI3K-mTOR — have been implicated in ASD (Takumi et al., 2020; Marlin and Froemke, 2017; Quezada, 2011). Key molecules within this latter pathway, such as PTEN, play critical roles in regulating cell differentiation, proliferation, and apoptosis (Rademacher and Eickholt, 2019).

Although genetics play a significant etiological role in ASD (Chaste and Leboyer, 2012), it is important to consider the influence of environmental factors. One such factor is Chlorpyrifos (CPF), an organophosphate pesticide, which has been linked to ASD in preclinical and human studies (Biosca-Brull et al., 2021). Gestational CPF exposure induces various adverse effects, including malformations, developmental delays, and functional impairments (Guardia-Escote, 2021). Furthermore, several studies have demonstrated deficits in innate and learned sociability and fewer ultrasonic vocalizations (USV) following CPF exposure (Lan et al., 2017; Lan et al., 2019; Morales-Navas et al., 2020).

ASD is also commonly associated with gastrointestinal issues, metabolic deficits, and dysbiosis (Tuchman, 2013; Vuong and Hsiao, 2017). Recently, the microbiota-gut-brain axis has gained attention as a potential etiological factor in ASD. In germ-free models, alterations in sociability have been observed in paradigms such as the Crawley test, where reduced motivation to interact with conspecifics or respond to social novelty is evident (Crumeyrolle-Arias et al., 2014). Microbiota can modulate host behavior and brain function by directly activating the vagus nerve, promoting inflammation, or stimulating metabolite production (Sampson and Mazmanian, 2015). For example, *Firmicutes* (e.g., *Lactobacillus*) and *Bacteroidetes* can modulate or directly produce precursors of vitamins and neurotransmitters (e.g., GABA, glutamate, serotonin), as well as bile acids, cytokines and short-chain fatty acids such as butyrate and acetate (Peralta-Marzal et al. 2021; Sampson and Mazmanian, 2015; Carabotti et al., 2015). Consequently, there has been growing interest within the scientific community in identifying effective gestational probiotic treatments, as probiotics can impact offspring development. This can occur either through bacterial colonization of the fetus via the placenta and amniotic fluid or by influencing fetal development through changes in maternal health (Swartwout and Luo, 2018).

Probiotic supplementation, particularly with strains of *Lactobacillus* and *Bifidobacterium*, has been shown to improve autistic symptoms in humans and mitigate ASD-like behaviors in preclinical studies (Shaaban et al., 2018; Sunand et al., 2020; Fung et al., 2017; Hsiao et al., 2013; Vuong and Hsiao, 2017). These empirical findings have led to the increased commercialization and usage of such agents, often combined with other compounds such as vitamin D (VitD) (Battistini et al., 2020). Together, probiotics and VitD have proven beneficial in treating depression and ASD, regulating the expression of central tyrosine hydroxylase, and reducing oxidative stress and inflammation (Abboud et al., 2020; Guiducci et al., 2022). Moreover, the gene encoding the vitamin D receptor (VDR) is one of the few genes that has “survived” rigorous genetic meta-analysis in ASD research (Mpoulimari and Zintzaras, 2022). Prenatal Vitamin D treatment has also been shown to restore sociability and learning impairments in ASD models (Vuillermot et al., 2017; Siracusano et al., 2020).

Therefore, based on the established link between prenatal CPF exposure and ASD, our study aims to replicate the effects of this exposure in offspring (both male and female). Additionally, we seek to investigate the safety and efficacy of probiotic supplementation combined with VitD as a potential treatment, assessing whether these alterations can be reversed. It was hypothesized that prenatal CPF exposure would induce an ASD-like phenotype, observable through behavioral tasks assessing social and communicative behaviors, as well as through changes in gut metabolites, brain gene expression, or epigenetic markers. Furthermore, probiotic and Vitamin D supplementation is expected to block or reduce these neurotoxic events at both behavioral and molecular levels.

## 2. METHODOLOGY

### 2.1 Experimental animals

The study sample consisted of 158 Wistar rats of both sexes, bred in-house at our facilities. Progenitors were obtained from Envigo (France). On postnatal day (PND) 1 (with birth designated as PND0), all pups were sexed, separated from their original mothers, and randomly assigned to foster mothers within the same experimental group. Each foster mother received 10 pups, with an equal distribution of sexes (half females and half males) to ensure a representative population and minimize bias related to genetic background. Weaning occurred on PND21 (see Figure 1), and thereafter, animals of the same sex and group were housed together in groups of four per cage. The animal housing room was maintained at a constant temperature of 22 ± 2 °C and humidity of 50 ± 10%, with a 12-hour light/dark cycle (lights on at 19:00h). Rats had *ad libitum* access to standard diet (SAFE®) and water throughout the study. The experimental protocol was conducted following the Spanish Royal Decree 53/2013 and the European Community Directive (2010/63/EU) on animal research and approved by the University of Almeria Animal Research Committee. Furthermore, the study adhered to the ARRIVE guidelines.

**Figure 1.**
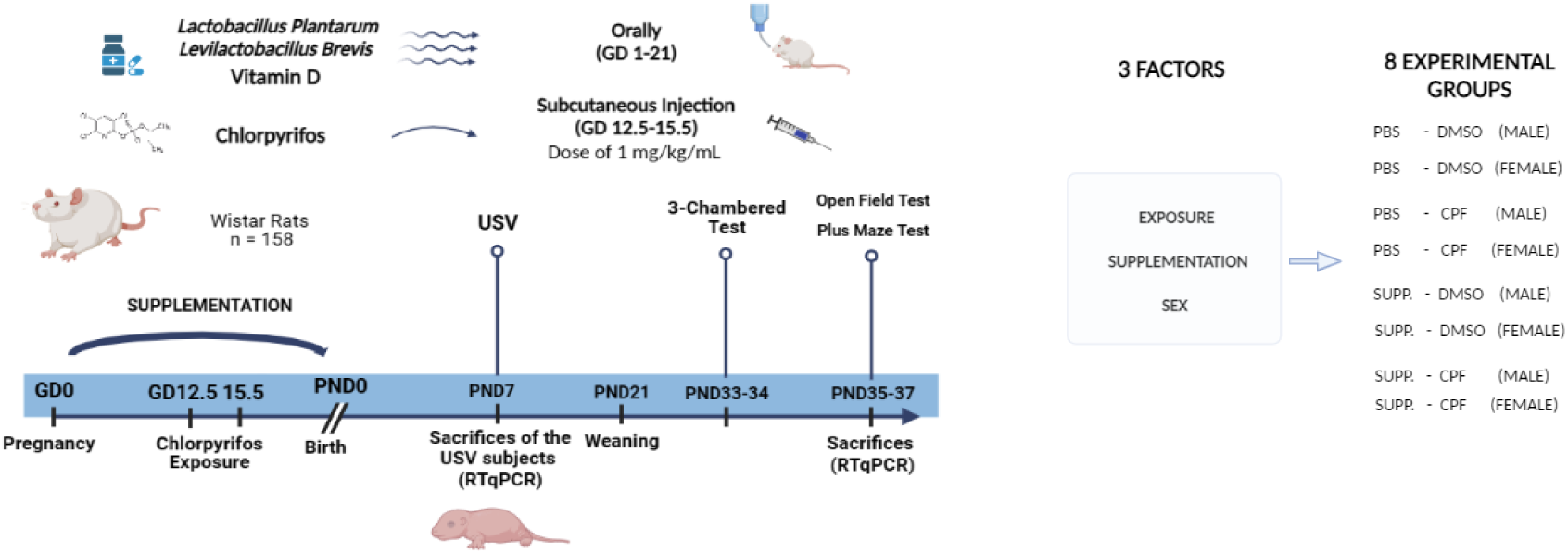
Experimental design. Timeline created using Biorender®.

### 2.2 Neurotoxic Agent

CPF [O, O-diethyl O-3,5,6-trichloropyridin-2-yl phosphorothioate (Pestanal, Sigma Aldrich)] was administered to pregnant dams via subcutaneous injection from gestational day (GD) 12.5 to 15.5, between 12.00 and 13:30h. Half of the animals were randomly assigned to the CPF exposure group, while the remaining animals were exposed to the vehicle (DMSO). This gestational window was chosen based on previous research linking CPF exposure during this developmental stage to ASD-like traits in rodents (Morales-Navas et al., 2021; Lan et al., 2017; Lan et al., 2019; Biosca-Brull et al., 2021). The CPF dosage was set at 1 mg/kg/mL per day, dissolved in DMSO (1 mL/kg), as this solvent can effectively dilute a wide variety of otherwise poorly soluble molecules (Galvao et al., 2014). In the control condition, DMSO was administered using the same method and volume as in the CPF group.

### 2.3 Probiotic and Vitamin D Supplementation

The probiotic treatment consisted of active strains of *Lactobacillus Plantarum* (KABP 023, CECT 7485) and *Levilactobacillus Brevis* (KABP 052, CECT 7480). These strains were chosen because *Lactobacillus* species have been observed to be more sensitive to CPF than other bacterial species (Condette et al., 2015). This supplementation was generously provided by AB BIOTICS and contained a concentration of 2e+11 cfu/g of active strains (ensuring that each subject received 2e+9 cfu/g). In addition, the supplementation contained 0.01 mg of Vitamin D3 and excipients, including 1623 mg glucose, 266.5 mg maltodextrin, 10 mg Silicon Dioxide (E-551), and 0.5 mg fructose. During pregnancy, dams received oral treatment via bottles equipped with safety balls to minimize waste. The dietary supplementation (SUPP) was diluted in 10 mL of PBS from GD1 until GD21. Dams in the control group had access to PBS at the same volume and using the same method. This design resulted in four groups based on exposure and supplementation conditions: (PBS-DMSO (control), PBS-CPF, SUPP-DMSO, and SUPP-CPF.

### 2.4 Evolution of Body Weight and Functional Battery (PND1-30)

The pups’ body weight, eye-opening, and neuromotor development were assessed. All tests were conducted in the animal housing room between 9 and 12 a.m. under dim lighting, with the same temperature and humidity conditions as previously described. Body weight measurements were taken at PND1, 5, 10, 15, 21 and 30. Eye-opening was evaluated daily from PND12 to15, with ocular aperture scored as follows: 0 (both eyes closed), 1 (one eye open), and 2 (both eyes open). These direct scores were then converted into percentages (0, 50, and 100%). Neuromotor development was assessed on PND16 using a battery of tests. Grip strength was evaluated by dragging the pup across a grid and assessing its resistance, scored as 0 (low resistance ability), 1 (some resistance to displacement), and 2 (strong resistance). Adherence capacity was then measured by placing the pup on a 60° inclined slope for 15 seconds, scored as 0 (inability to hold on), 1 (some difficulty), and 2 (fully developed ability). Climbing ability was then tested on the same slope and divided into three sections. The rat was placed at the bottom, and its progress was scored as follows: 0 (unable to climb beyond the first section), 1 (able to reach the intermediate section), and 2 (able to climb to the final section).

### 2.5 Ultrasonic vocalizations (USV)

At PND 7, half the pups from each group were randomly selected and separated from their dams 15 minutes before the test. They were moved to a room close to the experimental room, along with their respective litters, and covered with cotton wool to minimize sudden temperature changes. For USV recording, each pup was isolated for 5 minutes in a soundproof chamber (80×60×70 cm). A microphone (Dodotronic ultramic 250k) positioned approximately 10 cm from the animal captured the vocalizations. The USVs were processed using SeaWave v2.0 software (CIBRA) in a 16-bit, 250 kHz format. To avoid potential time-related biases, the recording order was counterbalanced across treatment conditions: the control group was tested first, followed by the CPF exposure group. Room humidity and temperature were maintained as previously described. Dim lighting was used during the pre-experimental phase, while isolation and recording took place in complete darkness. The USV assessments were carried out between 9 and 12 am. The DeepSqueak program, supported by a regional convolutional neural network (Faster-RCNN) and trained by the authors, was used for call detection and analysis. USVs ranging between 30 and 65 kHz were manually selected, as the literature has shown this to be the frequency range at which rat pups modulate calls in this type of isolation situation (Portfors, 2007). The isolation and USV recording were conducted on PND 7, a period identified as the optimal peak for call emission in rats (Wöhr and Schwarting, 2008; Hofer et al., 2001). The parameters analyzed using the DeepSqueak software included the total number of calls, calls per minute, latency to the first call, mean principal frequency (kHz), and principal frequency peak (kHz).

### 2.6 Three-Chambered Crawley test

The Crawley test consists of a central chamber and two equally sized chambers on each side (30×98×50 cm), separated by plexiglass walls with doors. This test was conducted during adolescence (PND33-34). During the habituation phase, the experimental rats were placed in the central chamber, and their motor behavior was monitored for 5 minutes. Subsequently, an unfamiliar rat, Stranger 1 (S1), was introduced into a wire mesh cage (23×15×23 cm) positioned in one of the side chambers. Once the habituation phase ended, the previously closed doors were opened, and the exploratory behavior of the experimental rat was recorded and analyzed for 10 minutes. In the third phase, a second unfamiliar rat, Stranger 2 (S2), was introduced into another wire mesh cage placed in the opposite side chamber, while S1 remained in its original position. The experimental rat continued exploring the apparatus for an additional 10 minutes. Locomotor activity was monitored throughout all phases. The behavior of the animals was recorded using Ethovision v3.1 software. The chambers were cleaned with 70% ethanol between subjects, and dim lighting was used during the test. For motor behavior analysis, the following parameters were recorded: total distance traveled (cm), time in motion (sec), mobility (frequency), velocity (cm/sec), immobility (sec), immobility (frequency), and strong mobility (sec). “Strong mobility” refers to instances where the rat moved very quickly (exceeding 95% of its previous total mobility). Regarding social behavior, two measures were taken: the time spent by the rat in each chamber and the time spent sniffing each animal (sec). The social interaction index (SI) was calculated as (S1-empty)/(S1+empty), comparing the time that experimental animals spent in the social chamber/sniffing Stranger 1 to the time they spent in the empty chamber/sniffing the empty cage during the second phase. Similarly, the social novelty reaction index (SNI) was calculated as (S2-S1)/(S2+S1). These indices were calculated using both chamber time and sniffing time. To assess social efficiency in each phase, a ratio was calculated between the time spent in the chamber and the time spent engaging in prosocial behavior (sniffing). The test was conducted between 9.00 and 12.00 am, under dim lighting and the same temperature and humidity conditions described earlier.

### 2.7 Open Field Test

Locomotor activity was evaluated in 8 plexiglass activity cages (39x39x15 cm) one day after analyzing sociability (PND35). Photocell technology connected to a computer was used to record the rats’ movements. The test was conducted between 9.00 and 12.00 am under dim lighting, with the same temperature and humidity conditions described previously. Behavior was automatically recorded using VersaMax®, and the data were collected with VersaData® software (PLC Control System SL). The rats were taken to the experimental room and allowed to habituate for one hour before the test (which lasted for 30 minutes). The rats were randomly assigned to different experimental cages, each cleaned with 70% ethanol before use. The variables measured included total distance traveled (cm), vertical activity (frequency of rearing), time in motion (sec), time spent in the center and at the margins of the cage (sec), and velocity (cm/sec).

### 2.8 Plus Maze Test

Anxiety was assessed one day after locomotor activity recording (PND36) using the Plus maze test. This apparatus, positioned 90 cm from the floor, consists of two open arms and two closed arms connected by a central platform. Rats were brought to the experimental room and allowed a 1-hour habituation phase. The test began with the rat placed on the central platform, after which they were allowed to freely explore the maze for 5 minutes. Ethovision v3.1 (Noldus) was used to track behavior. The dependent variables measured included entries into closed and open arms (frequency), total distance traveled (cm), time spent in open and closed arms (sec), mobility time (sec) and frequency, strong mobility time (sec) and frequency, immobility time (sec) and frequency, and velocity (cm/sec). An anxiety index (AI) was also calculated as the ratio of time spent in the open arms to the time spent in the closed arms. Between trials, the apparatus was cleaned with 70% ethanol, and the order of subjects was counterbalanced across different conditions. The same lighting, temperature, and schedule conditions were maintained as in the previous behavioral tests.

### 2.9 Molecular Analysis

#### 2.9.1 Sacrifice Protocol

The sacrifice protocol involved the euthanasia of 24 animals (12 females and 12 males) in equal numbers from each of the eight experimental groups (n = 3/group). The animals were euthanized two hours after completing the USV test on PND7 and sacrificed by fast decapitation, followed by immediate dissection of the brain hemispheres. Brain tissue samples were stored in RNase-free tubes (1.5 mL) and flash-frozen at −80°C to avoid RNA degradation until further use. All materials used for tissue extraction were either autoclaved or treated with ZAP (Invitrogen). The same sacrifice protocol was applied to adolescent rats after the behavioral tests (PND37), with one modification: the rats were first anesthetized with isoflurane to minimize suffering. Subsequently, a fresh brain dissection was performed to obtain frontal lobe and hypothalamus samples. Gut samples (PND7 rats) and feces samples (PND37 rats) were also collected and frozen at −80 °C before analysis.

#### 2.9.2 Brain Gene Expression: Reverse Transcription Quantitative Polymerase Chain Reaction (RTqPCR)

Samples were treated using Trizol reagent (Ambion) to isolate the RNA. The total RNA concentration for each sample was measured using Qubit fluorometry (Invitrogen). All samples were then normalized to a concentration of 100 ng/uL for reverse transcription into cDNA (Maxima first strand, Fisher Scientific). The resulting cDNA was then diluted in RNase-free water (factor 1:10) and used for the qPCR reactions. Each reaction contained SYBR green master mix, RNase-free water, primers for the genes of interest, and cDNA. Reactions were performed in duplicate, using a thermal cycler (StepOne v2.2.2, Applied Biosystems). The cycle threshold (Ct) values and melting curves were analyzed to identify abnormal patterns. Primers targeting exon sections of the gapdh gene served as housekeeping genes, while primers targeting intron sections were used to detect possible genomic contamination of the samples.

Efforts were made to ensure the absence of genomic contamination, very early expression ratios of the genes (< 30 Ct), or abnormalities in melting curves (e.g., unspecified peaks). Primer efficiency was confirmed using a serial 1:10 dilution factor for optimal amplification. Ct values for each gene were normalized to those of the Housekeeping gene (ΔCt) and the sample with the lowest expression, which served as an internal control (ΔΔCt). These ΔΔCt values were then converted to fold changes, as presented in the results section. On PND7, expression levels of 34 genes related to major neurotransmitter systems, ion channels, proinflammatory cytokines, and other biomolecular pathways were analyzed. In adolescence (PND37), 13 genes of interest in the frontal cortex and eight in the hypothalamus were examined. Primer sequences are listed in Appendix 6.

#### 2.9.3 Epigenetic RTqPCR (microRNA)

The same samples used for mRNA were also used for microRNA (miRNA) analysis with the Applied Biosystems TaqMan MicroRNA Assays. Samples were normalized to a concentration of 10 ng/uL, as our pilot studies revealed that this concentration provided optimal expression levels. Each TaqMan assay involved a cDNA retrotranscription specific to each microRNA, using a master mix (composed of dNTPs, H2O RNAase-free water, RNAase inhibitors, and RT buffer), cDNA sample, and primers (5x). Retrotranscription was conducted in a thermal cycler, and the resulting product was immediately used for RTqPCR. For RTqPCR, each reaction included the TAQMAN master mix, RNAase-free water, primers (20x), and the retrotranscription resulting from each sample. Data analysis followed the same procedures used for mRNA, except that U6 snRNA served as a housekeeping gene. miRNA-124a and miRNA-132 (ThermoFisher) were selected for analysis due to their demonstrated importance in autism, sociability, and neurodevelopmental processes (Anitha and Thanseem, 2015).

### 2.10. NMR-based metabolomics experiments

#### 2.10.1. Sample extraction

Samples were freeze-dried for 72 hours. For both PND7 and PND37 rats, 20 mg of freeze-dried intestines and fecal matter, respectively, were extracted using 700 µL of a 1:1 mixture of CH_3_OH-d_4_ and D_2_O KH_2_PO_4_ buffer 1.5M (pH 7.4), containing sodium 3-(trimethylsilyl) propionic-2,2,3,3-d_4_ acid (TSP, 0.1%, w/v) and sodium azide (NaN_3_, 90 µM) as an enzyme inhibitor. The samples were sonicated for 20 minutes, stirred at 600 rpm for 10 minutes, and centrifuged at 13500 rpm for 5 minutes. The supernatants were then transferred to 5 mm NMR tubes.

#### 2.10.2. NMR Acquisition, Bucketing Metabolite Assignment, and Quantification

A Bruker Avance III 600 MHz spectrometer, operating at 600.13 MHz and equipped with a thermostatted SampleJet autosampler (500 positions) and a 5 mm QCI quadruple resonance pulse field gradient cryoprobe, was employed to conduct the ^1^H NMR experiments. ^1^H NMR spectra were acquired at 300 ± 0.1 K without rotation using the Bruker pulse sequence cpmgpr1d. The acquisition parameters were as follows: 32 scans, 16 dummy scans, 32K data points, a spectral width of 22.0 ppm, an acquisition time of 1.24 s, a relaxation delay of 3.0 s, FID resolution of 0.81 Hz, and a total echo timeLof 96Lms. Locking was performed on the CH_3_OH-d_4_ frequency. Metabolite assignments for both matrices were based on data derived from 2D NMR experiments (homo- and heteronuclear), corresponding to ^1^H−^1^H COSY, ^1^H−^1^H TOCSY, ^1^H−^13^C HSQC, and ^1^H−^13^C HMBC, supplemented by NMR databases (Chenomx and HMDB) and relevant literature. Quantification of metabolites was performed relative to the internal standard (TSP) by integrating peak areas from isolated signals. For the PND37 group, female and male cohorts were analyzed separately due to sex-specific metabolic differences in fecal samples.

### 2.11 Statistical analysis

Frequentist analyses were conducted using IBM SPSS Statistics 25, with three independent factors, each with two levels: exposure (DMSO or CPF), supplementation (PBS or SUPP), and sex (male or female). A repeated measures analysis of variance (ANOVA) was conducted to evaluate body weight changes and variables across blocks in the open field test, considering the three factors mentioned above. Additionally, a three-way ANOVA was used to analyze the functional battery, ocular opening, total number of USVs, plus maze test variables, and sociability variables recorded on the Crawley test. Gene and epigenetic expression analyses were based on fold change (relative expression) values using an ANOVA for each mRNA or miRNA of interest. Molecular outcomes are presented as individual plots in the figures, with p values provided in the text. Statistical significance was set at p < 0.05. The figures in the results section (mean ± SEM or individual values) display only significant interaction effects, excluding the sex factor due to its lack of significant effects in the analyses. GraphPad Prism v10.1.0 was utilized to create the figures.

For the metabolomics data, all NMR spectra were bucketed by selecting individual peaks in the region from δ_H_ 0.2 to 10.0 ppm using AMIX 3.9.15 (Bruker BioSpin GmbH, Rheinstetten, Germany). Buckets were then obtained by integrating the corresponding spectral areas. The δ_H_ 4.90−4.70 ppm and δ_H_ 3.31-3.34 ppm regions, which contain residual H2O suppression and methanol signals, were excluded from bucketing and subsequent analysis. Before conducting statistical analyses, the data were normalized by scaling the intensity of individual peaks to the total intensity of the entire spectrum, excluding the regions mentioned above. Multivariate data analysis was conducted using SIMCA-P software (v. 17.0, Umetrics). Both exploratory unsupervised analyses, such as PCA, and supervised models, such as PLS-DA, were applied to the ^1^H NMR data with scaling to unit variance. PLS-DA models were validated by assessing the goodness-of-fit (R^2^) and goodness-of-prediction (Q^2^) cumulative values, along with the CV-ANOVA parameter validation (at the significance level of p < 0.05) to evaluate model accuracy. Discriminant peaks were identified from loadings obtained from predictive models where the variable importance projection (VIP) plot exceeded 1. Fold change (FC) for discriminant metabolites with p < 0.05 (according to ANOVA) was analyzed using Fisher’s Least Significant Difference (LSD) post hoc test in Metaboanalyst. Bar plots were generated using GraphPad Prism v10.1.0.

## 3. RESULTS

### 3.1 Supplementation Advances General Development and Increases USV emissions, while CPF Exposure Blocks USV Increases

All variables related to postnatal development are summarized in Fig.2. Body weight evolution during the postnatal period (PND1 to PND30) revealed a significant SUPP*PND interaction effect [F(1.3,101.9)= 9.258, p < 0.001]. Significant differences in body weight were observed on PND5, 10, 15, 21, and 30 (p= 0.004, p= 0.008, p < 0.001, p= 0.001, p= 0.001, respectively),with the SUPP groups showing higher weights than their controls, regardless of CPF exposure (Fig.2A). For ocular opening analysis (Fig.2B), a significant SUPP*PND interaction was again detected [F(1,79)= 5.115, p= 0.026]. Post-hoc analysis revealed that the SUPP group showed a higher percentage of ocular opening than controls on PND13 (p= 0.003), but this difference disappeared by PND14 (p= 0.519). Analysis of the functional battery data revealed a significant effect of SUPP for both grip capacity [F(1,79)= 8.107, p= 0.006] and climbing capacity [F(1,79)= 10.460, p= 0.002]. The SUPP group demonstrated earlier motor development than controls (p= 0.006, p= 0.002), regardless of gestational CPF exposure (Fig.2C, Fig.2D). No significant differences were found for adherence capacity (Appendix1A).

**Figure 2.**
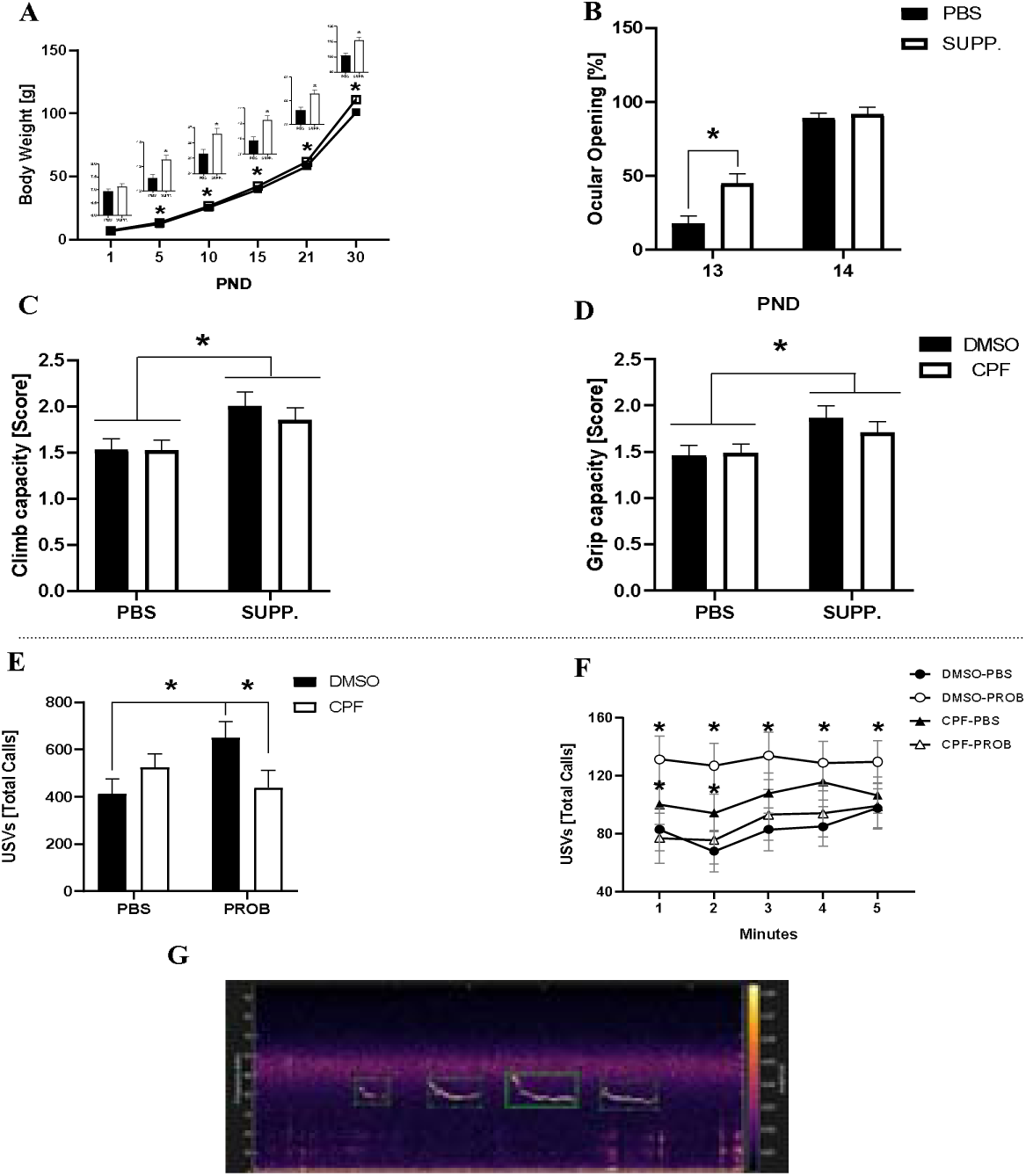
General development. Sample size: PBS-DMSO-MALE (n= 9), PBS-DMSO-FEMALE (n= 15), PBS-CPF-MALE (n= 15), PBS-CPF-FEMALE (n= 12), SUPP-DMSO-MALE (n= 10), SUPP-DMSO-FEMALE (n= 6), SUPP-CPF-MALE (n= 8), SUPP-CPF-FEMALE (n= 12). Mean (±SEM) of body weight evolution (A), percentage of ocular opening on PND13-14 (B), and scores obtained on the functional battery for grip capacity (C) and climbing capacity (D) are depicted. Additionally, the mean (±SEM) of the total number of calls (USV) emitted by the different groups (E) and the mean of the total number of USVs per minute (F) are represented. Sample size: PBS-DMSO-MALE (n= 8), PBS-DMSO-FEMALE (n= 10), PBS-CPF-MALE (n= 11), PBS-CPF-FEMALE (n= 10), SUPP-DMSO-MALE (n= 7), SUPP-DMSO-FEMALE (n= 8), SUPP-CPF-MALE (n= 6), SUPP-CPF-FEMALE (n= 7). *p < 0.05. The image provided in Figure G shows an example of USV from the DeepSqueak program for a single subject randomly selected to illustrate the software and its detection quality.

Regarding the data for USV emissions (PND7), as depicted in Fig.2, we observed an EXPOSURE*SUPP interaction for the total number of calls emitted by the pups during isolation (Fig.2E) [F(1,59)= 6.171, p= 0.016]. Under normal conditions, SUPP significantly increased the number of calls (p= 0.012), an effect completely blocked by CPF exposure (p= 0.037).

Subsequently, a more detailed analysis was conducted on the evolution of the number of USVs emitted per minute (Fig.2F). This interaction was only significant only during the first 4 minutes of isolation [F(1,59)= 5.338, p= 0.024; F(1,59)= 6.755, p= 0.012; F(1,59)= 4.417, p= 0.040; F(1,59)= 5.276, p= 0.025]. Throughout the session (at least statistically during the first 4 minutes), the overall increase in USVs observed was always compared to the control condition (p= 0.030, p= 0.007, p= 0.025, p= 0.033, respectively). However, the ability of CPF exposure to significantly reduce calls in the SUPP group was noted only during the first two minutes (p= 0.025, p= 0.028). Regarding behavioral topography, no effects were observed for the analyzed variables (latency, mean frequency, average peak frequency, or mean USV duration). These non-significant results are presented in Appendix 1B.

### 3.2 Supplementation Enhanced Spontaneous Exploratory Activity in Response to a Novel Environment, With These Differences Disappearing Following Habituation

A significant SUPP*BLOCK interaction was found for both total distance traveled [F(5,355)= 5.878, p < 0.001] and mobility time [F(5,355)= 4.922, p< 0.001]. SUPP increased the total distance traveled in the first 3 blocks (p= 0.007, p < 0.001, p= 0.041, respectively), although these effects disappeared in the last 3 blocks of the test (p= 0.271, p= 0.953, p= 0.315) (Fig. 3A). A similar pattern was observed for the time spent by the animals in motion (Fig. 3B), where SUPP groups remained significantly longer in motion than controls during the first 3 blocks (p= 0.009, p<0.001, p= 0.041), with these effects again disappearing in the last 3 blocks (p= 0.429, p= 0.804, p= 0.294). All these effects were independent of CPF exposure. Analysis of rearing behavior (Fig.3C) revealed the significant differences underlying the SUPP*BLOCK interaction [F(5,355)= 3.660, p= 0.003]. SUPP increased the frequency of vertical activity in Blocks 1 (p= 0.008) and 2 (p= 0.024). No significant differences were found for anxiety-related variables, the time spent in the margins, or time spent in the center of the apparatus (Appendix.2A). Finally, CPF exposure in both SUPP and PBS groups increased total velocity in the OFT [F(1,70)= 4.868, p= 0.031] as shown in Appendix 2A.

**Figure 3.**
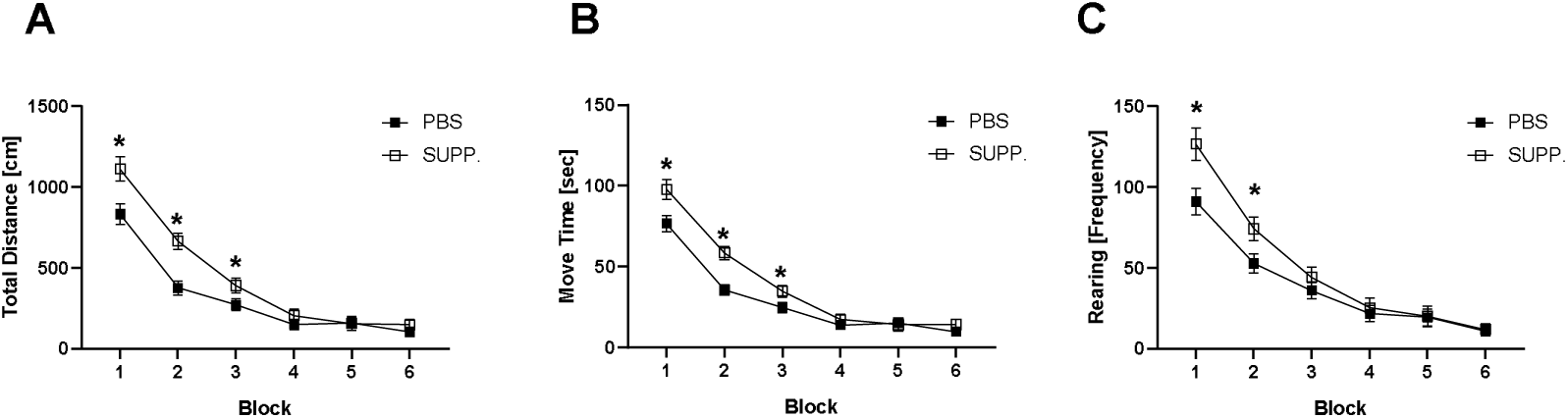
Locomotor activity in the OFT. Mean (±SEM) total distance traveled (A), time that animals spent in motion (B), and rearing frequency (C) in each 5-minute block. Sample size: PBS-DMSO-MALE (n= 10), PBS-DMSO-FEMALE (n= 10), PBS-CPF-MALE (n= 15), PBS-CPF-FEMALE (n= 12), SUPP-DMSO-MALE (n= 10), SUPP-DMSO-FEMALE (n= 5), SUPP-CPF-MALE (n= 7), SUPP-CPF-FEMALE (n= 11). *p < 0.05

### 3.3 Prenatal CPF Reduces Anxiety Indicators During Adolescence, Both Under Normal Conditions and in Supplemented Rats

While no alterations in anxiety were observed in the OFT, in the plus maze, prenatal CPF exposure significantly reduced the anxiety index (Fig.4A) regardless of SUPP administration[F(1,71)= 9.400; p= 0.003]. This reduction is attributed to CPF increasing the time spent in the open arms (Fig.4B) [F(1,71)= 9.239; p= 0.003] and decreasing the time spent in the closed arms (Fig.4C) [F(1,71)= 6.325; p= 0.014]. Although supplementation did not directly affect anxiety levels in our sample, a significant EXPOSURE*SUPP interaction was observed [F(1, 71)= 5.026, p= 0.028] for immobility frequency (Fig.4D). Under normal conditions, probiotic and VitD gestational supplementation increased the number of times the rat stopped (p< 0.001), while CPF blocked this effect (p= 0.417), differing from the SUPP group (p= 0.006). However, no significant differences were found in the duration of immobility (p= 0.121).

**Figure 4.**
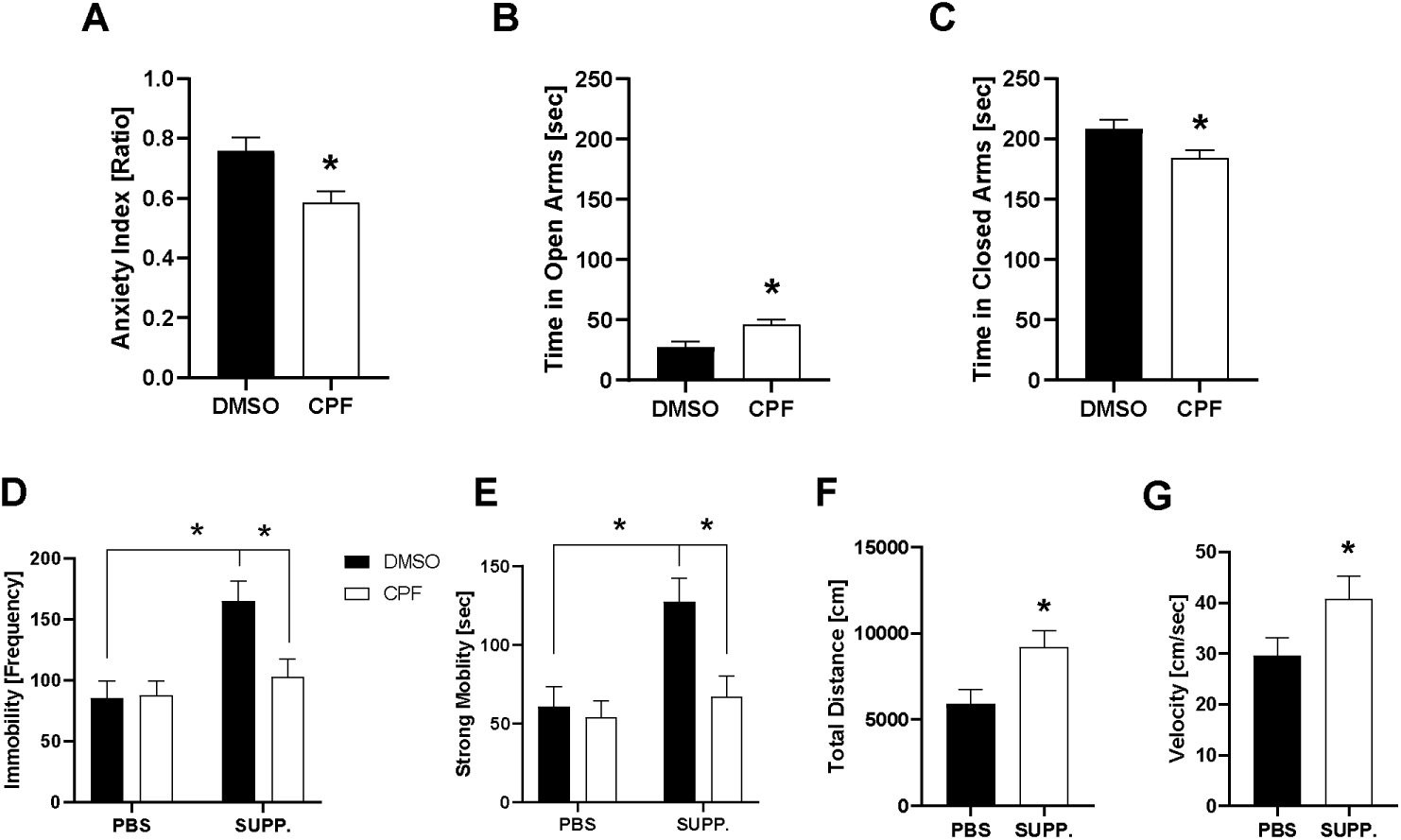
Elevated Plus Maze Test. Means (±SEM) are shown for the anxiety index (A), time spent in the open arms (B), closed arms (C), frequency of immobility (D), time spent performing strong mobility (E), total distance traveled (F), and velocity (G). Sample size: PBS-DMSO-MALE (n= 10), PBS-DMSO-FEMALE (n= 10), PBS-CPF-MALE (n= 15), PBS-CPF-FEMALE (n= 12), SUPP-DMSO-MALE (n= 10), SUPP-DMSO-FEMALE (n= 5), SUPP-CPF-MALE (n= 7), SUPP-CPF-FEMALE (n= 11). *p<0.05

While no differences were observed in mobility frequency and duration (see Appendix 2B), the same pattern of results was replicated for the frequency of strong mobility (Fig.4E) [F(1,71)= 4.231, p= 0.043]. In this case, supplementation increased the number of instances of sudden and heavy mobility (p= 0.001), while the CPF group did not differ significantly from the control group (p= 0.443) and exhibited significantly less of this behavior than the SUPP group (p= 0.004). Moreover, total distance traveled (Fig.4F) and velocity (Fig.4G) were significantly increased by supplementation, independent of CPF exposure [F(1,71)= 6.522, p= 0.013; F(1,71)= 3.966, p= 0.05; respectively].

### 3.4. Supplementation Failed to Restore Social Behavior Deficits in CPF-Exposed Rats While Altering Responses Social Novelty in the Control Group

In the sociability phase (S1 vs. Empty Cage), a significant EXPOSURE*SUPP interaction [F(1,59)= 5.095, p= 0.028] was found for the time spent in the social chamber (Fig.5A). CPF exposure decreased this time only in the SUPP group (p= 0.028). However, no significant differences were found in the time spent interacting with S1 [F(1,55)= 0.766, p= 0.385] (Fig.5C). In contrast, CPF increased the time animals spent in the chamber without a social stimulus (Fig.5B) [F(1,59)=7.671, p= 0.007]. This finding was supported by the SI data, which revealed that CPF decreased the percentage of time spent in the chamber with S1 compared to the empty chamber [F(1,59)= 5.712, p= 0.020] (Appendix 3A). The same pattern of results was observed when SI (Fig.5D) was calculated based on sniffing time [F(1,55)= 4.815, p= 0.032]. Furthermore, CPF exposure significantly reduced the percentage of time spent engaged in sniffing behavior, indicating lower social efficiency [F(1,55)= 4.140, p= 0.047] (Fig.5E). In the social novelty reaction phase (S1 vs S2), CPF exposure increased the time the animals spent in the room with S2 (Fig.5G) [F(1,59)= 7.981, p= 0.006]. However, no significant increase was observed in sniffing toward the novel animal due to CPF, indicating that this increase may not correspond to prosocial behavior toward social novelty. Nonetheless, supplementation led to a decrease in sniffing toward S2 [F(1,56)= 4.560, p= 0.037] (Fig.5H). On the other hand, supplementation significantly increased the time spent in the room with the familiar animal (S1) [F(1,59)= 9.253, p= 0.003]. Post-hoc analysis revealed an EXPOSURE*SUPP interaction [F(1,59)= 13.458, p= 0.001] where the increase induced by SUPP was blocked by CPF exposure (p= 0.004) (Fig.5F). These increase (p= 0.028) and blocking (p= 0.042) effects were also observed for sniffing toward S1 [F(1,56)= 5.598, p= 0.021], suggesting a rise in prosocial behavior toward the familiar animal (Appendix 3A). These effects can be further understood by examining the effect on the SNI, which showed that supplementation decreased this value for both the time chamber ratio [F(1,59)= 7.369, p= 0.009] and the time sniffing ratio [F(1,54)= 6.136, p= 0.016]. The EXPOSURE*SUPP interaction (Fig.5I) [F(1,54)= 5.607, p= 0.021] revealed that the SUPP group showed a clear preference for the familiar animal or S1 (p= 0.002), which was blocked by prenatal exposure to CPF (p= 0.013). In this case, no statistically significant differences were observed in social efficiency (Fig.5H). Outcomes for motor variables are presented in Appendix 3B.

**Figure 5.**
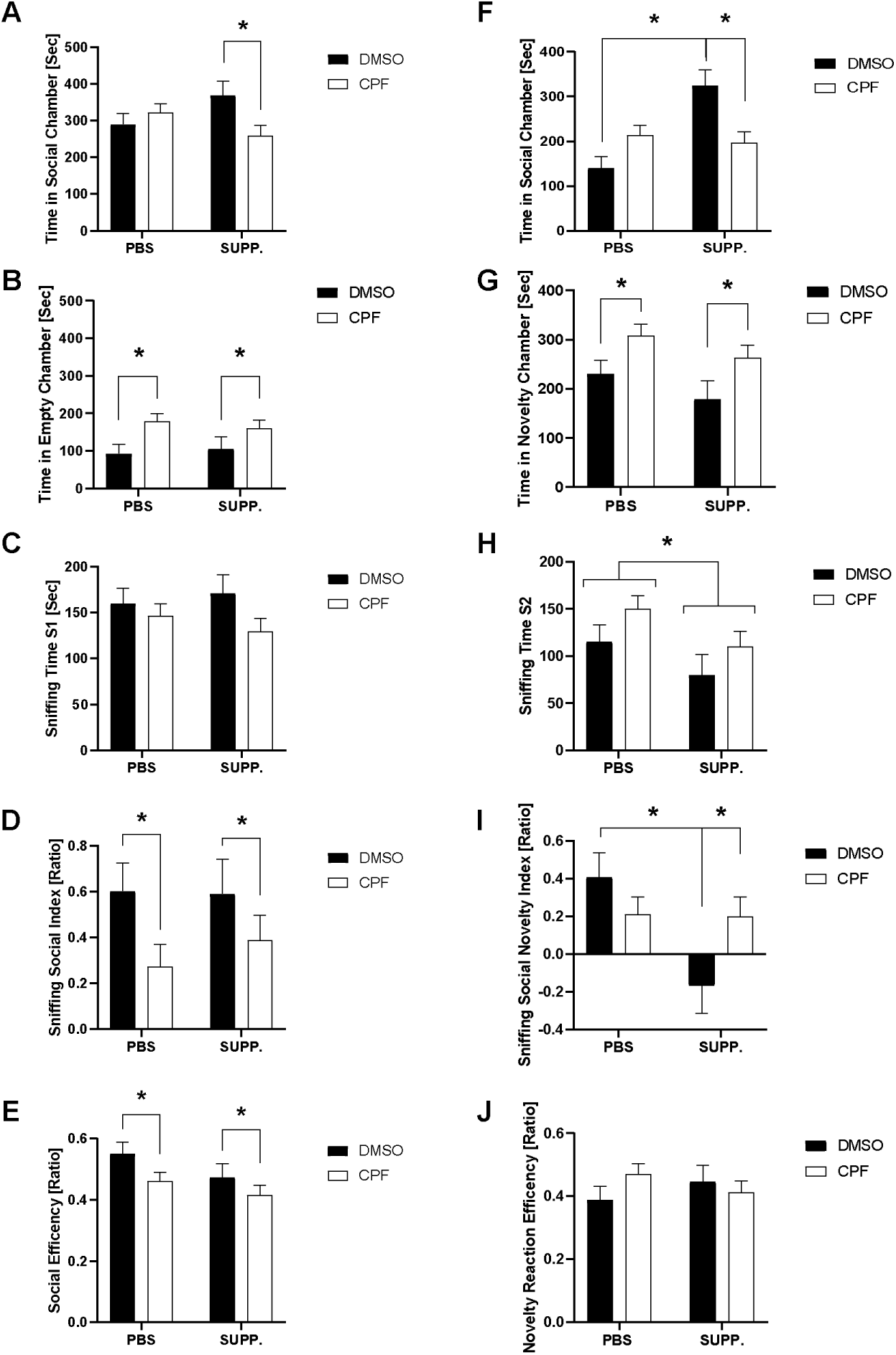
3-Chambered Test. For the sociability phase, mean (±SEM) values are depicted for various parameters: Time spent by the animal in the social room with S1 (A), and in the empty room (B), time spent engaged in sniffing as a prosocial behavior with S1 (C), the social index calculated using sniffing time (D), and social efficiency, defined as the proportion of time the animal spent engaged in prosocial sniffing behavior with S1 relative to the total time spent in the social room(E). Additionally, the following social novelty reaction phase variables are also represented: Time spent by the animal in the familiar room with S1 (F) and time spent by the animal in the chamber with the novel social stimulus, S2 (G), time spent sniffing in response to the unfamiliar social stimulus, S2 (H), the social novelty index calculated from sniffing time (I) and the efficiency in social novelty reaction (J). Sample size: PBS-DMSO-MALE (n= 6), PBS-DMSO-FEMALE (n= 10), PBS-CPF-MALE (n= 12), PBS-CPF-FEMALE (n= 10), SUPP-DMSO-MALE (n= 8), SUPP-DMSO-FEMALE (n= 3), SUPP-CPF-MALE (n= 8), SUPP-CPF-FEMALE (n= 10). *p < 0.05 indicates statistical significance.

### 3.5. Supplementation Altered the Expression of ASD-Related Genes in the Pups’ Brains, with CPF Restoring These Alterations to Normal Levels

Analysis of the expression of genes related to social behavior in the entire cerebral hemisphere, illustrated in Fig.6, revealed an EXPOSURE*SUPP interaction for both sexes on the oxytocin receptor (*Oxyr*) and the vasopressin receptor subfamily 1a (*Avpr1a*) [F(1,16)= 6.521, p= 0.025; F(1,16)= 5.391, p= 0.034; respectively]. Analysis of this interaction revealed that supplementation increased the expression of these genes (p= 0.011, p= 0.047), while CPF blocked this effect (p= 0.034, p= 0.006). However, for *Avpr1a* expression, we also observed a general downregulation effect due to gestational CPF exposure, irrespective of supplementation [F(1,16)= 4.629, p= 0.047]. Moreover, the same EXPOSRE*SUPP interaction effect was found for other ASD-related genes: *Pten* [F(1,16)= 9.332, p= 0.008], *Maoa* [F(1,16)= 7.897, p= 0.013], and *Foxp1* [F(1,16)= 4.786, p= 0.044], where supplementation increased their expression (p= 0.022, p= 0.008, p= 0.049), while brief CPF exposure abolished these effects in the co-exposed group (p= 0.012, p= 0.007, p= 0.029). The same interaction effect was also found for the expression of *Kcc2* and the α2 subunit of the GABA-A receptor (*Gabra2*) [F(1,16)= 5.684, p= 0.030; F(1,16)= 9.519, p= 0.007], with supplementation increasing their expression (p= 0.043, p= 0.044), and CPF blocking the effect (p= 0.018, p= 0.016). For *Pi3kr1* expression (Appendix 4), this interaction [F(1,16)= 7.734, p= 0.013] was solely due to differences in the SUPP group, where CPF reduced its expression (p= 0.025). Finally, supplementation, regardless of CPF exposure, led to a downregulation of NMDA receptor subunit 2c (*Grin2c*) [F(1,16)= 4.984, p= 0.040]. Appendix 4 presents the genes for which no statistically significant differences were found.

**Figure 6.**
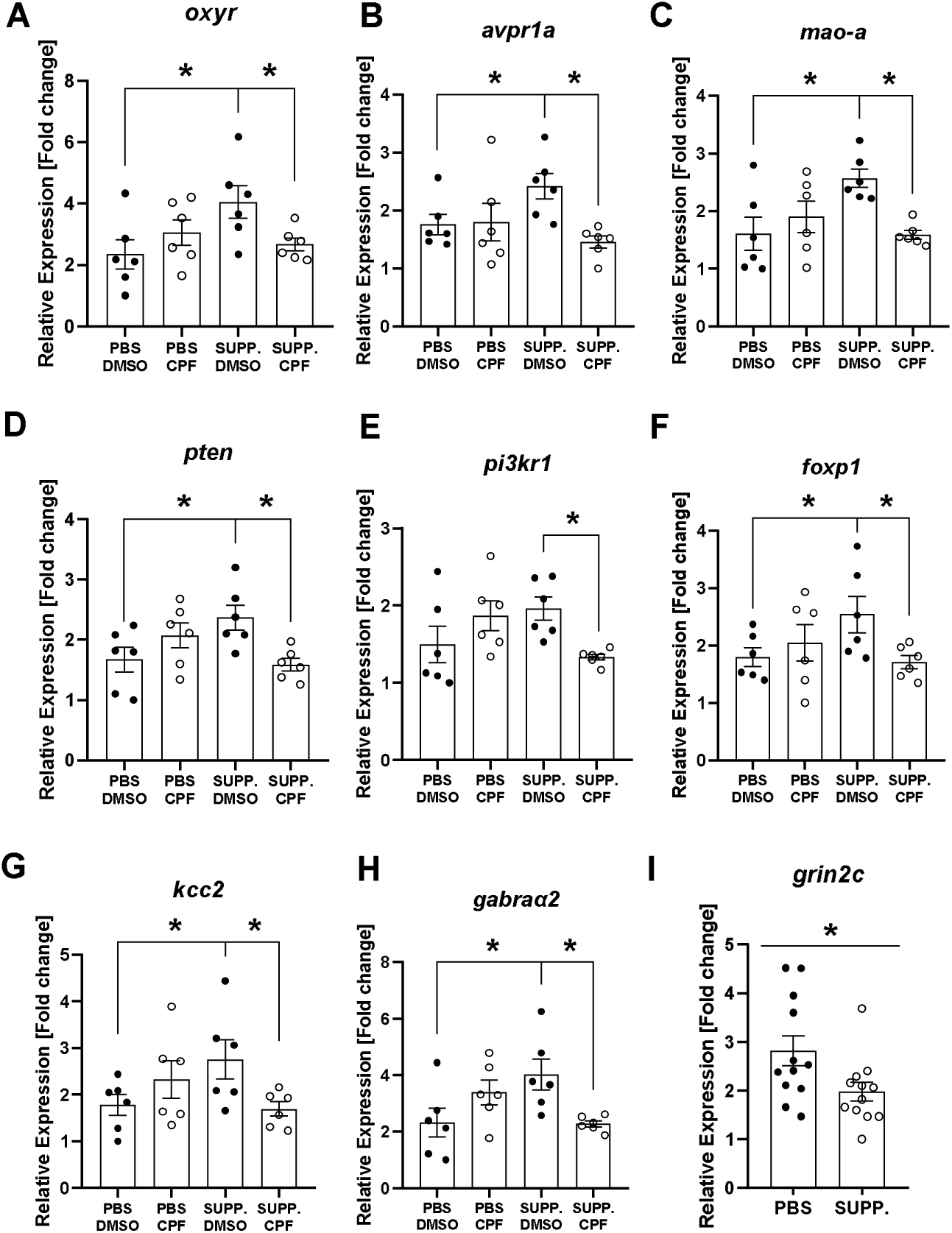
Gene expression (mRNA). The mean (±SEM) gene expression in the whole cerebral hemisphere during infancy (PND7) are: oxytocin receptor (A), vasopressin receptor subfamily 1a (B), monoamine oxidase A enzyme (C), phosphatase and tensin homolog (D), phosphoinositide-3-Kinase Regulatory Subunit 1 (E), forkhead-box protein P1 (F), the chloride-potassium cotransporter KCC2 (G), α2 subunit of GABA-A receptor (H), and the SUPP-exclusive effect on NMDA receptor subunit 2c RNA levels (I).

### 3.6. CPF Exposure Increased miRNA-124a Levels in the Whole Hemisphere of the Offspring, an Effect Counteracted by the Dietary Supplementation

As shown in Fig.7, at PND7, supplementation did not affect the whole cerebral hemisphere with respect to *miRNA-124a* [F(1,16)= 0.011, p= 0.918] or miRNA-132 [F(1,16)= 1.164, p= 0.297]. However, we observed an effect on *miRNA-124a* in the EXPOSURE*SUPP interaction [F(1,16)= 6.033, p= 0.026], where CPF increased the expression of *miRNA-124a* (p= 0.050), an effect that was mitigated by supplementation, restoring *miRNA-124a* levels to those observed in the control group (p= 0.195).

**Figure 7.**
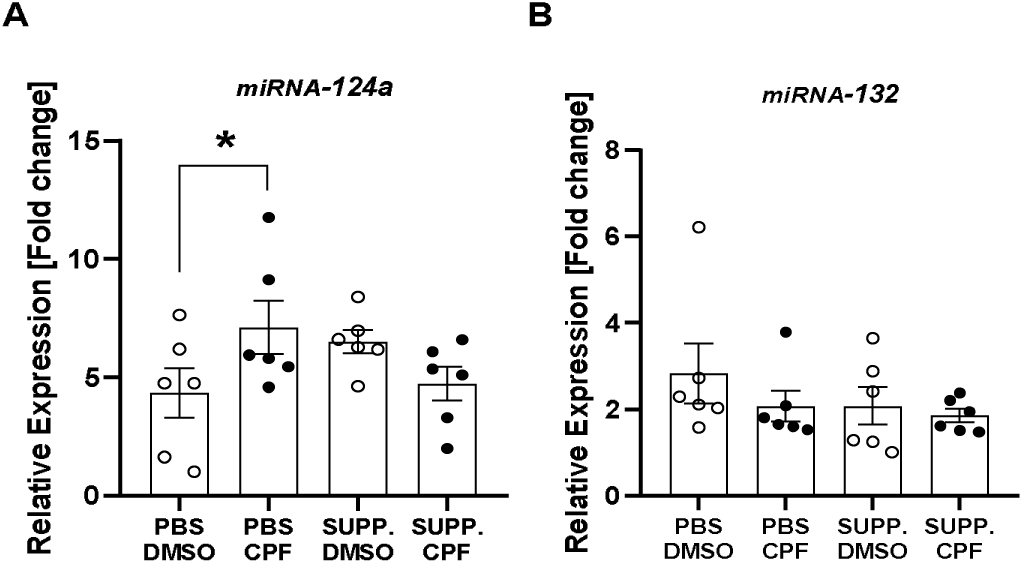
Epigenetic (miRNA). Mean(±SEM) levels of *miRNA-124a* (A) and *miRNA-132* (B) in the whole brain hemispheres. *p> 0.05

### 3.7. Gene Expression Pattern Changes Observed in the Frontal Cortex and Hypothalamus During Adolescence in *Avpr1a* and *Kcc2*

The results of RTqPCR in adolescence are presented in Fig.8. In the frontal cortex, a significant effect on gene expression of Avpr1a was found in the EXPOSURE*SUPP interaction [F(1,16)= 4.717, p= 0.045]. However, in this case, the expression pattern was opposite to that found during early development. Supplementation produced a down-regulation of *Avpr1a* (p= 0.037), which was reversed by CPF exposure (p= 0.035). The same interaction was observed for *Kcc2* [F(1,16)= 8.568, p= 0.010], where the CPF-PBS group showed higher levels of expression in the frontal cortex (p= 0.040), which was corrected by supplementation (p= 0.009). No statistically significant differences were found for *Nkcc1, Kcc1, Maoa, Pten, Oxyr*, or any of the RNA levels in the analyzed subunits of the GABA-A receptor (α1, α2, β2, and γ2), *Grm,* or 2c NMDA receptor subunit (*Grin2c*) (See Appendix 5). In the hypothalamus of the same subjects, we observed a significant increase in the expression of *Avpr1a* as a result of supplementation [F(1,16)= 13.612, p= 0.002], regardless of CPF exposure. Significant differences were also observed in the expression of *Kcc2*. Specifically, CPF strongly down-regulated this cotransporter in the hypothalamus [F(1,16)= 29.877, p< 0.001], while supplementation induced an increase in its expression in this area [F(1,16)= 5.370, p= 0.034]. No significant differences were found for *Nkcc1, Oxyr, Gabra2, Grin2c,* or *Pten* (Appendix 5).

**Figure 8.**
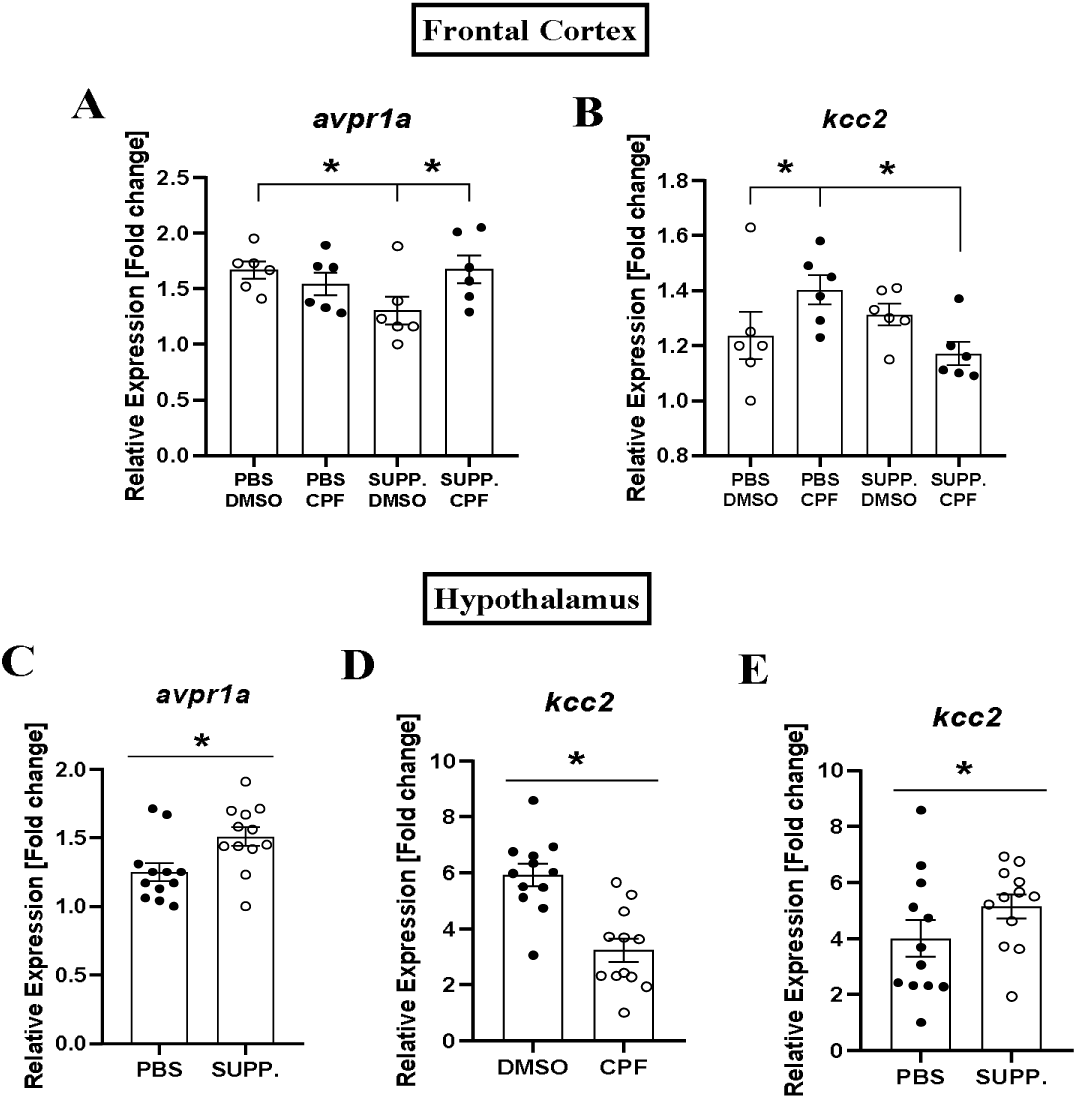
Gene expression (mRNA) During Adolescence. Mean relative gene expression levels (±SEM) of *Avpr1a* (A, C) and *Kcc2* (B, D, E) in frontal cortex and hypothalamus are depicted.

### 3.8. Significant Alterations in Gut and Fecal Metabolites Following Supplementation and CPF Exposure

Metabolite assignments were conducted by NMR and are detailed in Tables S1 and S2, respectively. A total of 38 metabolites were identified across guts and feces, including bile acids, major short-chain fatty acids (SCFAs) produced by the intestinal microbiota (acetic, propionic, butyric acid, and 3-hydroxybutyric acid) (Hiseni et al., 2023), carbohydrates (glucose, galactose, sucrose), amino acids (valine, isoleucine, leucine, alanine, lysine, proline, arginine, glutamate, glutamine, aspartate, glycine, and phenylalanine), intermediates of glycine

metabolism (sarcosine and N, N-dimethylglycine), quaternary ammonium compounds such as choline, and a choline-derivative tentatively assigned to glycerophosphocholine (Wang et al., 2005) and betaine, along with their precursor ethanolamine, and several organic acids (acetate, lactate, fumarate, and formate). Regarding gut metabolites on PND7, a valid PLS-DA model was established between the SUPP and PBS groups exposed to CPF, considering both male and female samples (Fig.9A). Discrimination was primarily observed in several spectral regions, all with a variable importance in projection (VIP) value exceeding 1. These regions are highlighted in orange in the loadings plot displayed in Fig.9B. Relative peak integrals of discriminant metabolites from these regions with VIP > 1 in the PLS-DA model and with p < 0.05 based on ANOVA with Fisher’s Least Significant Difference (LSD) post hoc test are depicted in Fig.9C.

**Figure 9.**
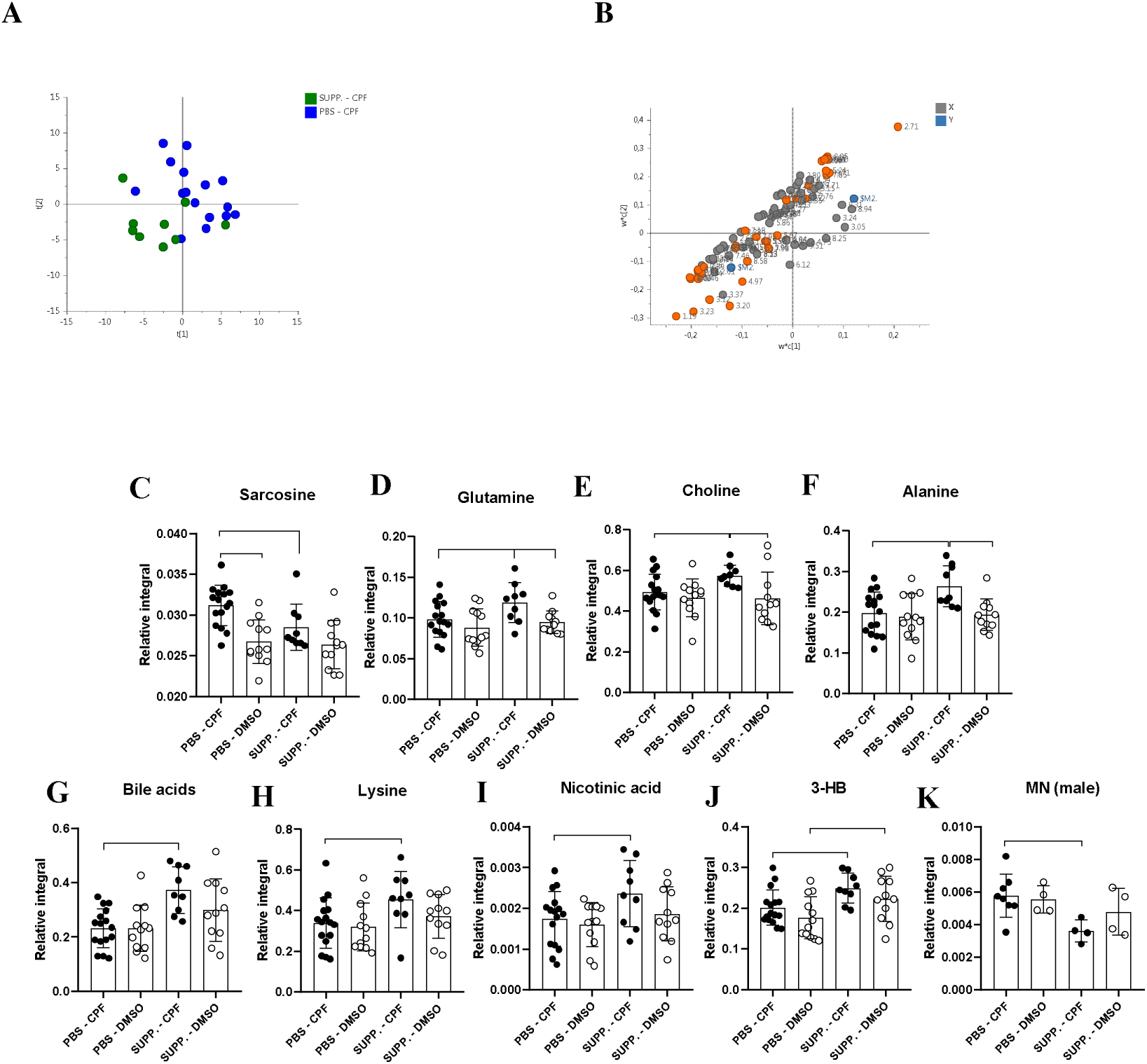
PLS-DA scores plot of ^1^H NMR data from guts at PND7 in rats of both sexes belonging to SUPP and PBS groups and those exposed to CPF (scaling to UV, R^2^ = 0.72, Q^2^ = 0.55, CV-ANOVA =0.048) (A). PLS-DA loadings plot highlighting, in orange, the discriminatory spectral regions between the two groups with a variable importance in projection (VIP) > 1 (B). Relative peak integrals of discriminant metabolites with VIP > 1 in PLS-DA model and with p < 0.05 according to ANOVA followed by Fisher’s LSD test. Means ±SEM are depicted for sarcosine (C), glutamine (D), choline (E), alanine (F), bile acids (G), lysine (H), nicotinic acid (I) and 3-hydroxybutyrate (J) considering both sexes, except for 1-methylnicotinate (K), which was only statistically discriminant in males. Sample size: PBS-DMSO-MALE (n= 6), PBS-DMSO-FEMALE (n= 9), PBS-CPF-MALE (n= 11), PBS-CPF-FEMALE (n= 8), SUPP-DMSO-MALE (n= 6), SUPP-DMSO-FEMALE (n= 7), SUPP-CPF-MALE (n= 6), SUPP-CPF-FEMALE (n= 7).

Sarcosine is the N-methyl derivative of glycine and is an intermediate in the metabolic pathway of glycine. This metabolite significantly increased in the CPF-PBS group compared to the others (p = 0.0001, p(FDR) = 0.012), indicating that CPF exposure increased sarcosine levels, an effect blocked by supplementation. Additionally, increased levels of the amino acids glutamine (FC= 1.19, p = 0.025), alanine (FC= 1.33, p = 0.0030) and lysine (FC= 1.30, p = 0.027) were observed in this group, whereas choline (FC= 1.12, p = 0.049), nicotinic acid (FC= 1.25, p = 0.031) and bile acids (FC= 1.61, p = 0.0005) increased as a result of supplementation in the CPF-exposed group. Supplementation also decreased gut 1-methylnicotinate (trigonelline) levels of CPF-exposed male rats (FC = 0.62, p = 0.0082) but significantly increased short-chain fatty acid 3-HB levels regardless of CPF exposure in both sexes on PND7 (FC= 1.23, p = 0.019 and FC= 1.23, p = 0.022 for CPF and DMSO groups, respectively).

Additionally, regardless of supplementation, CPF exposure led to an increase in the following gut metabolites in both sexes (see Fig.10): betaine (FC= 1.26, p = 0.0016 and FC= 1.28, p = 0.035 for probiotic and PBS groups, respectively), ethanolamine (FC= 1.29, p < 0.0001 and FC= 1.24, p < 0.0001), N, N-dimethylglycine (FC= 1.14, p = 0.003 and FC= 1.13, p = 0.029) and taurine (FC= 1.12, p = 0.036 and FC= 1.26, p = 0.0001).

**Figure 10.**
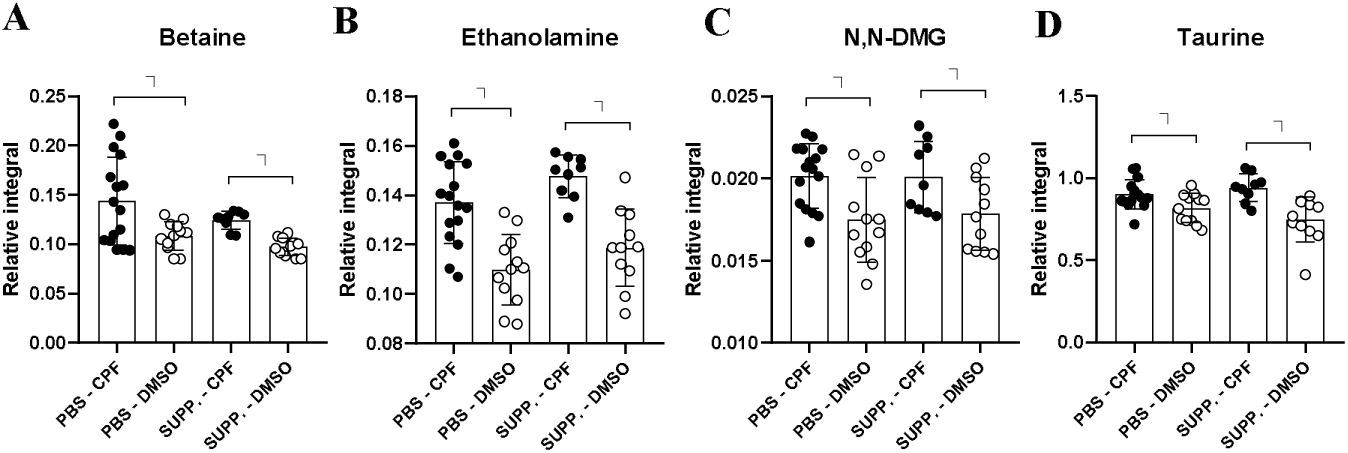
Bar plots depicting relative peak integrals of metabolites that increase in CPF-exposed groups regardless of supplementation in PND7 rats (p < 0.05 according to ANOVA followed by Fisher’s LSD test). Mean ±SEM values are shown for betaine (A), ethanolamine (B), N, N-dimethylglycine (C) and taurine (D) considering both sexes. Sample size: The same as for Figure 9.

Regarding PND37 fecal samples, female CPF-exposed rats given supplementation exhibited a 2-fold reduction in fecal bile acid levels compared to the PBS group (p = 0.014). However, the opposite outcome was observed in male rats: when CPF was administered along with SUPP, a 1.6-fold increase in bile acid content was observed compared to the CPF-exposed group without SUPP intake (p = 0.014). Additionally, CPF-exposed males given supplementation displayed a decrease in fecal glycine (FC = 1.56, p = 0.011), sarcosine (similar to PND7 rats, FC = 1.40, p = 0.013), and sucrose (FC = 1.63, p = 0.020) in comparison to their non-supplemented counterparts (Fig.11).

**Figure 11.**
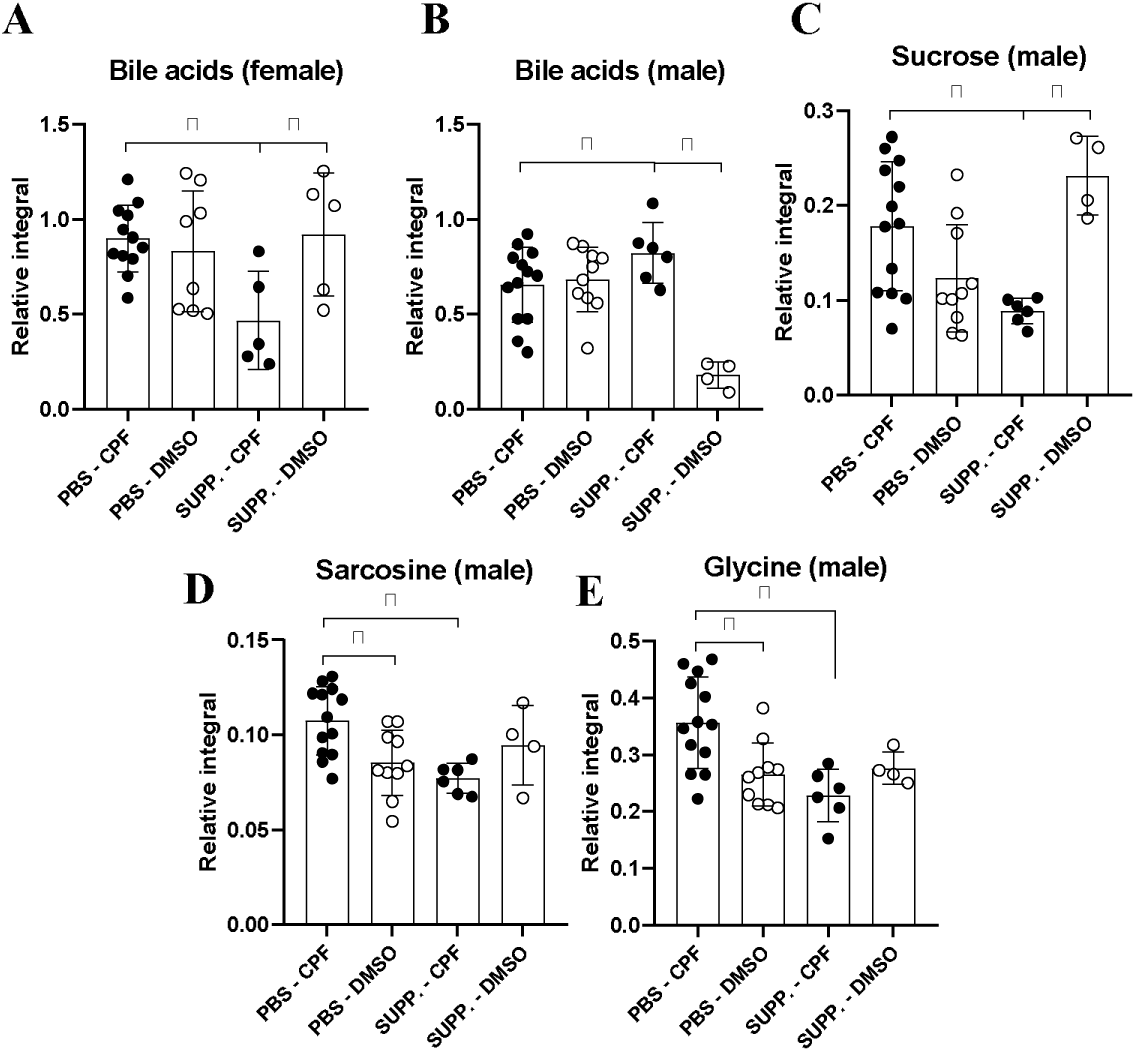
Bar plots depicting relative peak integrals of metabolites that differ between SUPP and PBS groups for CPF-exposed rats at PND37 (p < 0.05 according to ANOVA followed by Fisher’s LSD test). Mean ±SEM of bile acids in females (A) and males (B), sucrose in males (C), sarcosine (D) and glycine (E) in males are shown. Sample size: PBS-DMSO-MALE (n= 10), PBS-DMSO-FEMALE (n= 8), PBS-CPF-MALE (n= 13), PBS-CPF-FEMALE (n= 12), SUPP-DMSO- MALE (n= 4), SUPP-DMSO-FEMALE (n= 5), SUPP-CPF-MALE (n= 6), SUPP-CPF-FEMALE (n= 5).

Concerning CPF exposure effects, no significant differences were observed between female CPF-exposed and DMSO groups. Conversely, in male rats, higher levels of the amino acids glutamine (FC= 1.34, p= 0.003 and FC= 2.07, p < 0.0001 for SUPP and PBS groups, respectively), aspartate (FC= 1.31, p= 0.032 and FC= 1.55, p= 0.014) and proline (FC= 1.25, p=0.020 and FC= 2.31, p < 0.0001) and N, N-dimethylglycine (FC= 1.28, p= 0.017 and FC= 1.67, p= 0.0019) were observed for CPF-exposed groups compared to DMSO groups, regardless of supplementation (Fig.12).

**Figure 12.**
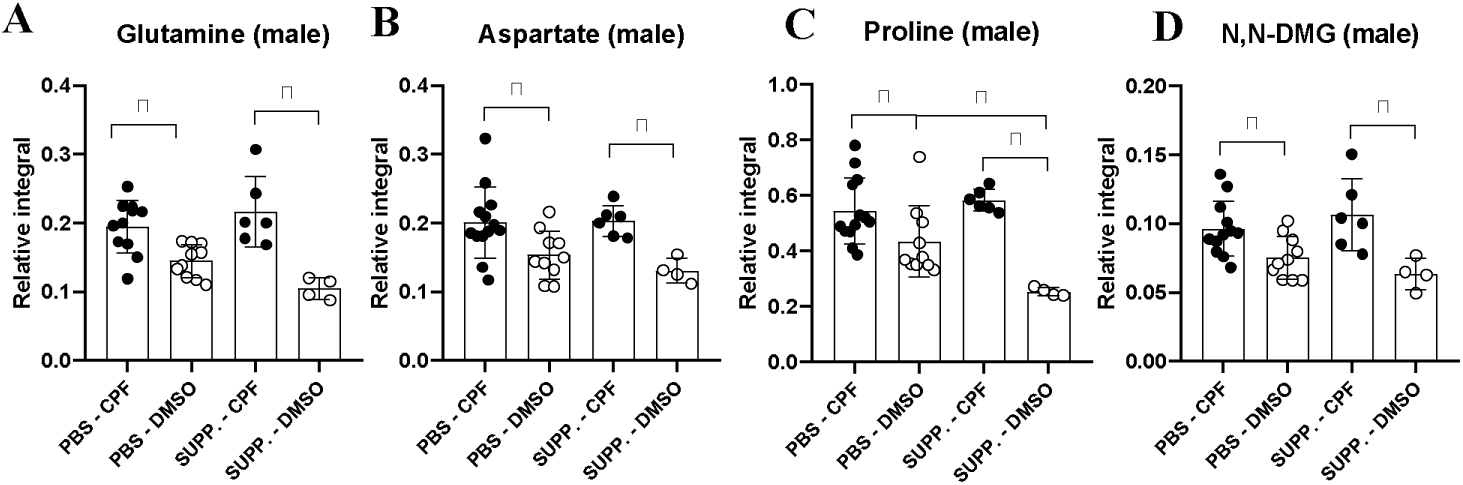
Bar plots depicting relative peak integrals of fecal metabolites that increase for CPF-exposed groups regardless of SUPP at PND37 (p < 0.05 according to ANOVA followed by Fisher’s LSD test). Mean ±SEM values are shown for glutamine (A), aspartate (B), proline (C) and N, N-dimethylglycine (D) in males. Sample size: The same as for Figure 11.

## 4. DISCUSSION

We anticipated that CPF exposure would induce ASD-like behaviors and alterations in gene expression commonly associated with this disorder and that supplementation would potentially reverse these effects. However, our findings indicate that supplementation can induce significant changes in development, behavior, brain gene expression, and metabolism in non-CPF-exposed controls. Surprisingly, CPF emerges as the agent capable of inhibiting these effects. Irrespective of whether the changes induced by supplementation are deemed positive or negative, we have demonstrated its significant impact on a control organism. Rats whose mothers were treated throughout gestation with a supplement composed of the bacterial strains *L. Plantarum* and *L. Brevis*, combined with Vitamin D, exhibited increased body weight gain, an advancement in general and motor development, and changes in several neurodevelopmental genes. This developmental advancement is accompanied by increased USV emissions and a potential strengthening of attachment bond formation, manifesting as a preference for the social stimulus to which the animal has been habituated instead of social novelty. Both behavioral effects appear to be blocked by prenatal CPF exposure, a pattern of results similar to that observed in brain gene expression. To the best of our knowledge, this is the first time these effects have been observed following this dietary supplementation in control rats. In adolescence, CPF exposure impairs sociability; however, supplementation does not appear to correct this alteration.

Perinatal administration of *L. Rhamnosus* has been shown to reduce excessive body weight gain (Luoto et al., 2010). However, prenatal administration of Vitamin D has been associated with increased body weight, while a deficit in prenatal Vitamin D has been linked to low birth weight in offspring in the short and medium term (Pérez-López et al., 2020). The low dose of glucose used as an excipient makes it unlikely to cause weight gain. Studies using higher doses, such as Li et al. (2023), have shown no weight difference in offspring, only reduced birth weight. The increase in body weight observed in our sample could be explained by an overall advancement in development, as evidenced by the greater percentage of subjects that exhibited earlier eye-opening and improved climbing and gripping abilities. Furthermore, supplementation resulted in an increase in brain gene expression of *Kcc2* on PND7. This increase, along with the upregulation of *Gabra2* and the downregulation of *Grin2c*, suggests a potential neurodevelopmental advancement in establishing the GABAergic system as inhibitory (Peerboom and Wierenga, 2021). Such advancement could have long-term effects on the formation of the E/I (excitatory/inhibitory) balance, which is often affected in ASD. Early advancement of *Kcc2* expression leads to a weakening of glutamatergic synapses, as certain developmental windows rely on the depolarizing function of GABA to form these synapses. Conversely, an immediate increase is observed in GABAergic transmission, which involves the α1 subunit (Peerboom and Wierenga, 2021; Huang et al., 2012) and possibly, as observed in our study, the α2 subunit. CPF exposure blocked this neurodevelopmental advancement.

Supplemented pups (at PND7) exhibited an increase in the number of USVs, an effect blocked by brief prenatal CPF exposure. This blocking effect partially aligns with the results reported by Morales-Navas et al. (2020). At the molecular level, the same pattern is found in gene expression in pups’ brains. Supplementation increased the mRNA levels encoding oxytocin receptors, the vasopressin receptor family 1a, *Maoa*, *Pten*, and *Foxp1* in the brain, an effect again blocked by CPF. Alterations in *Pten* and its influence on the PI3K/AKT/mTOR pathway can influence the expression of *Foxp1* (Garcia-Forn et al., 2020). This family of transcription factors has been associated with the development of issues in vocal communication and language learning (Garcia-Oscos et al., 2021). Additionally, MAO-A has been shown to interact with FOXP2, which is also associated with vocal communication (Park et al., 2014). It has been demonstrated that Vitamin D levels during early development regulate the amount of FOXP2 in the pup rat cortex (Yates et al., 2018). Furthermore, MAO-A is implicated in disorders associated with antisocial behavior (Davis et al., 2008; Park et al., 2014). Conversely, the oxytocin and vasopressin neurotransmission systems are involved in sociability and communication, systems also affected in ASD (Marlin and Froemke, 2017; Quezada, 2011).

Finally, at PND7, levels of the miRNAs of interest were examined to elucidate the potential post-transcriptional epigenetic mechanisms involved. *miRNA-132*, associated with neurodevelopment, synaptic plasticity, and inflammatory phenomena (Anitha and Thanseem, 2015), is altered in individuals with ASD and with exposure to medium-high doses of CPF (Lee et al., 2016). However, we found no such alterations in any of our groups. In contrast, *miRNA-124a*, which promotes neural differentiation and regulates spatial learning and social interactions (Anitha and Thanseem, 2015), increased due to CPF exposure. This is the first instance in which this miRNA has been shown to be affected by CPF exposure. The findings reported by Bahi (2016) indicate an overexpression of *miRNA-124* in the hippocampus related to reduced sociability as a consequence of social isolation in neonatal Wistar rats (PND1-11), leading to a decrease in the expression of its endogenous target, BDNF. In our study, we observed a similar trend in *Bdnf* expression. It is plausible that the context of social isolation to which we exposed the offspring (at PND7) during USV recording produces epigenetic changes via *miRNA-124a* and that prenatal CPF exposure generates an epigenetic vulnerability, which is corrected by supplementation. Future studies employing social isolation models should further investigate the presence of these epigenetic vulnerabilities related to CPF exposure and explore the potential protective effect of supplementation.

In the longer term, during adolescence, CPF decreased sociability in Phase 2 of the Crawley test, as evidenced by reduced time spent in the social room and decreased prosocial sniffing behavior. These behavioral changes are associated with alterations in the gene expression of *Avpr1a* and *Kcc2* in the frontal cortex and hypothalamus. Interestingly, in our study, supplementation did not restore sociability in CPF-exposed rats, contrary to findings in the literature where gestational probiotics (Wang et al., 2019) and postnatal (Du et al., 2017) or prenatal Vitamin D supplementation (Vuillermot et al., 2017) have been shown to correct sociability deficits in ASD models such as those induced by valproic acid or maternal immune activation. The sociability deficits observed cannot be attributed to locomotor deficits or anxiety, as CPF-exposed rats displayed reduced anxiety levels. However, in the social novelty phase, CPF did not appear to affect the social novelty-seeking behavior typically observed in these animals (Beery and Shambaugh, 2021), with CPF-exposed rats even showing a preference for the novel chamber. In contrast, supplementation induced a paradigm shift in control rats, leading to a loss of preference for social novelty and a marked preference for the known animal. At the physiological level, gestational supplementation increased the long-term expression of *Kcc2* and *Avpr1a* in the hypothalamus, where VDR partially colocalized with vasopressin (Liu et al., 2021). In the frontal cortex, however, supplementation corrected the CPF-induced increase in *Kcc2* expression and decreased *Avpr1a* expression. Vasopressin has been widely implicated in specific aspects of sociability, such as attachment bond formation, attachment preference responses to social novelty, and monogamy in both human and non-human animals (Walum et al., 2008; Phelps, 2010). This system, along with the *Avpr1a* gene, is implicated in ASD, and vasopressin administration has been shown to increase affiliative behavior toward conspecifics (Kim et al., 2002). Interestingly, CPF blocked the down-regulation of *Avpr1a* in the frontal cortex and the associated behavioral attachment effects.

Supplementation significantly increased the time spent in motion, the total distance covered in the first three 5-minute blocks of the OFT, and the rearing frequency. These findings suggest that subjects exhibited hyperactivity induced by exposure to a novel environment, with these effects gradually diminishing with habituation to the OFT. Similar results were reported by Liu et al. (2016), who used *L. Plantarum* postnatally in a germ-free rodent model during adolescence. This increase in activity occurred regardless of the anxiety levels in the environment, as evidenced by increased mobility and velocity in the elevated plus maze. Taken together, the evidence suggests that supplementation modulates the locomotor and exploratory activity of the host. Metabolomic analysis revealed that CPF exposure induced elevated levels of betaine, N, N-dimethylglycine, sarcosine (under normal conditions), and alanine (in the SUPP group). These findings suggest potential disruptions in the glycine metabolism pathway, which is crucial for anti-inflammatory responses and intestinal microbiota regulation (Wu et al., 2020).

Interestingly, imbalances in glycine are commonly observed in ASD (Abreu et al., 2021; Zheng et al., 2017; Smith et al., 2019). Additionally, the increase in gut levels of ethanolamine in CPF-exposed pups (SUPP group) and choline suggests potential disturbances in phosphatidylethanolamine and phosphatidylcholine synthesis pathways, which could impact methylation processes associated with ASD (Hamlin et al., 2013). Taurine, an amino acid known to increase in ASD (Ghanizadeh, 2013), was found to be elevated in the guts of CPF-exposed rats. Along with glycine, taurine can exist in free form and as a component of conjugated bile acids (Ijare et al., 2010). This increase in taurine in the SUPP-CPF group suggests potential alterations in specific gut bacteria related to bile acid metabolism, essential for lipid metabolism, maintaining intestinal barrier function, and regulating energy homeostasis (Zhao et al., 2016). Alterations in bile acid metabolism, as seen in the BTBR model of ASD, are often accompanied by changes in gut *Lactobacillus* populations and several other microbial taxa that produce bile salt hydrolase enzymes, facilitating the de-conjugation of bile acids from urine or glycine (Golubeva et al., 2017).

Additionally, an increase in nicotinate (the acid form of vitamin B3) was observed in the SUPP-CPF group, together with a decrease in its metabolized product, 1-methyl nicotinate. This compound plays a vital role in energy metabolism and possesses gut anti-inflammatory properties (Santoru et al., 2020), suggesting a positive effect of the supplementation on inflammation-driven metabolism caused by CPF. Moreover, 3-HB levels increased due to supplementation, independent of CPF exposure. This short-chain fatty acid, derived from microbial metabolism, is known for its various physiological activities, including the suppression of inflammatory diseases (Suzuki et al., 2023) and its protective effects against extracellular stress, as well as enhancing synaptic plasticity (Zhu et al., 2023). In adolescent fecal samples, comparable patterns in glycine and bile acid metabolism influenced by CPF exposure and supplementation were identified, revealing sexual dimorphism in bile acid levels.

More caution should be exercised regarding the indiscriminate use of dietary supplements, especially during pregnancy, as they appear to significantly influence the behavior and biology of control rats. Probiotics, often marketed as supplements rather than drugs, lack stringent post-market safety reporting requirements (Merenstein et al., 2023). Theoretically, it has been suggested that probiotics could produce harmful effects in hosts, such as systemic infections, alterations in neurodevelopment, changes in metabolic activity, excessive immune stimulation, impacts on metabolism, and modulation of drug and toxic effects (Merenstein et al., 2023; Doron and Snydman, 2015). Additionally, continuous high-dose Vitamin D treatment can be toxic (Battistini et al., 2020) and could potentially exacerbate autoimmune responses in the brain (Häusler et al., 2019). However, caution is warranted due to the inherent differences in intestinal microbiota and general physiological function between rats and humans, which make direct comparisons impossible. We propose the necessity for systematic and thorough safety testing of prenatal supplementations, given the physiological potential shown by the supplementation used in our study.

The primary limitation of this study is the inability to identify which component of the supplementation contributes to the phenotypes observed in the offspring. Future research should explore the mechanisms underlying the observed changes across various levels of analysis and attempt to distinguish which effects can be attributed to the probiotic component, the specific dose of Vitamin D, or any of the other excipients used, as well as how these components interact within the organism. The small amount of glucose (1.6 mg) administered, compared to the high concentration of glucose present in standard rat food, suggests that these substances are unlikely to significantly account for the neurodevelopmental outcomes observed. Moreover, previous studies using rats to investigate the effects of glucose consumption on inducing metabolic disorders and brain alterations have utilized much higher doses (Moreno-Fernández et al., 2018), supported by the absence of alterations in insulin or glucose metabolism in our study. In addition, the composition of the intestinal microbiota was not assessed in this study due to a focus on metabolite analyses and the lack of available samples. Further research is needed to investigate the changes in gut microbiota that accompany these effects. Finally, we did not include a pure control of the vehicles used to administer the treatments. Thus, future experiments would benefit from including such controls.

In conclusion, gestational supplementation with probiotics and Vitamin D unexpectedly produced stable, robust, and long-term effects in control rats. These effects included a general advancement in development, increased USV emissions during social isolation, a preference for familiar rats over social novelty, hypermotility, and increased exploratory behavior. Co-exposure with gestational CPF blocked several of these behavioral effects, suggesting involvement in the development of GABAergic, glutamatergic, oxytocinergic, and vasopressinergic systems regulating autism-like behaviors. Furthermore, our findings confirm the hypothesis that CPF significantly reduces sociability in offspring, rendering the supplementation ineffective in addressing these deficits. Thus, further research is needed to develop safer and more effective probiotic treatments.

## FUNDING AND ACKNOWLEDGEMENTS

This work was supported by the Spanish Government (Ministry of Science and Innovation; MCIN/AEI 10.13039/501100011033) (Reference: PID2020-113812RB-C32). Finally, we thank AB BIOTICS for providing the dietary supplementation used in this study.

## CRediT AUTHOR STATEMENT

**Coca, Mario R**: Investigation, Writing - Original draft preparation, Writing - review & editing, Visualization, Formal analysis, Conceptualization. **Abreu, Ana C.**: Writing - Original draft preparation, Investigation. **Salmerón Ana M**: Writing - Original draft preparation, Investigation. **Morales-Navas, Miguel**: Methodology, Software, Formal analysis, Investigation, Writing - Original draft preparation, Supervision. **Ruiz-Sobremazas, Diego**: Investigation, Writing - Original draft preparation. **Colomina, Teresa**: Writing - Original draft preparation, Supervision, Conceptualization. **Fernández, Ignacio:** Writing - Original draft preparation, Supervision, Conceptualization. **Perez-Fernandez, Cristian**: Investigation, Writing - Original draft preparation, Writing - Review & editing, Visualization, Supervision, Methodology, Conceptualization. **Sanchez-Santed, Fernando**: Conceptualization, Methodology, Resources, Writing - Original draft preparation, Writing - Review & editing, Supervision, Conceptualization, Methodology, Resources, Writing - Original draft preparation, Writing - Review & editing, Supervision, Project administration, Funding acquisition.

## DECLARATION OF GENERATIVE AI AND AI-ASSISTED TECHNOLOGIES IN THE WRITING PROCESS

The author(s) generated, reviewed, and edited the content as needed and take(s) full responsibility for the content of the publication. No generative AI or AI-assisted technologies were used in the writing process.

## DATA STATEMENT

The data supporting the findings of this study are available upon request. Researchers can access the data by contacting the corresponding author.

