## Appendix for "Modulating Neurotoxic Effects of Prenatal Chlorpyrifos Exposure Through Probiotic and Vitamin D Gestational Supplementation: Unexpected Effects on Neurodevelopment and Sociability"

**SUPPLEMENTARY MATERIAL**

**Appendix 1**

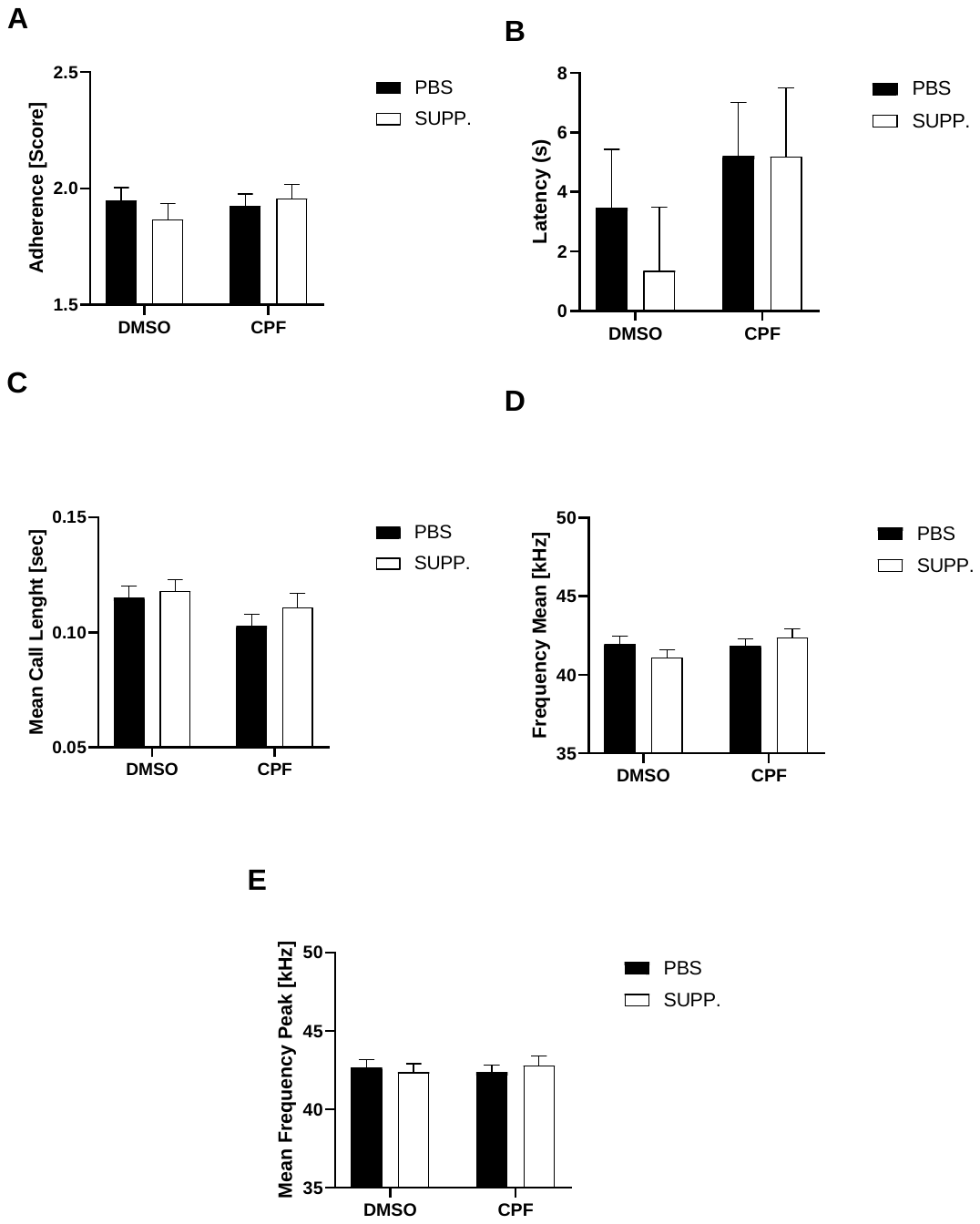

**Appendix 1.** Mean (±SEM) of adherence capacity on PND16 (A), USV emission latency (B), mean USV frequency (C), mean USV frequency peak (D) and the mean call length duration (E).

**Appendix 2A**

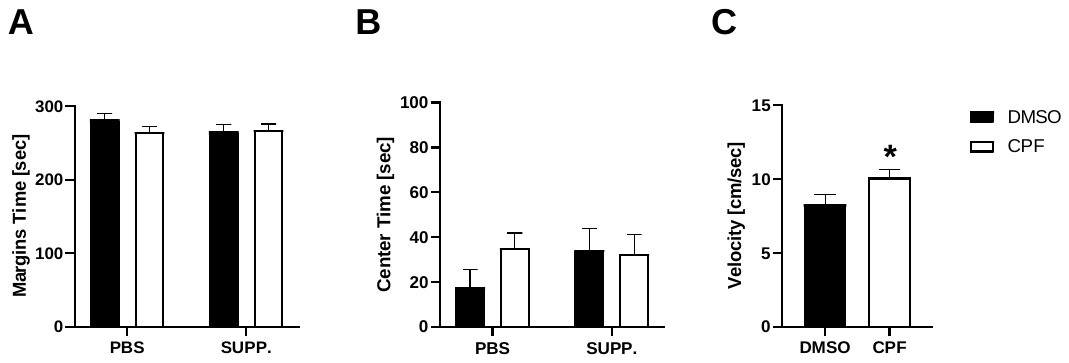

**Appendix 2A.** Open Field Test. Mean (±SEM) of time spent in margins (A) and center (B) of the paradigm, and mean velocity (C). *p< 0.05

**Appendix 2B**

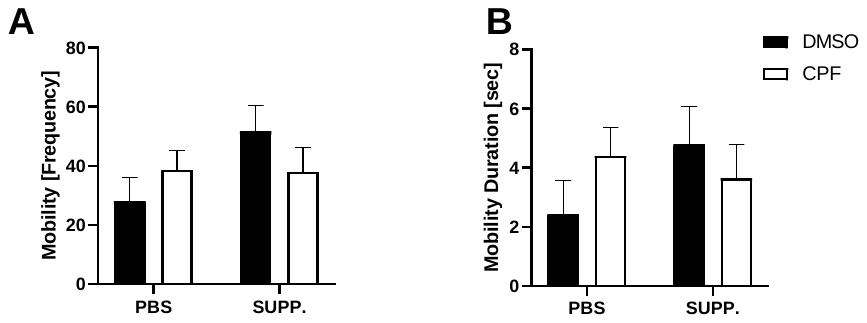

**Appendix 2B.** Elevated Plus Maze Test. Mean (±SEM) of mobility frequency (A) and duration (B).

**Appendix 3A**

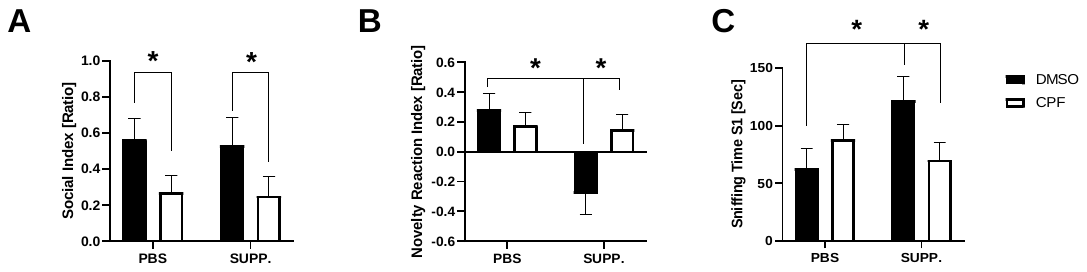

**Appendix 3A.** 3-Chambered Test. Mean (±SEM) of social (A) and novelty reaction index (B) calculated with time spent in chambers. Sniffing time with S1 (familiar animal) in 3 phase (C). *p< 0.05

**Appendix 3B**

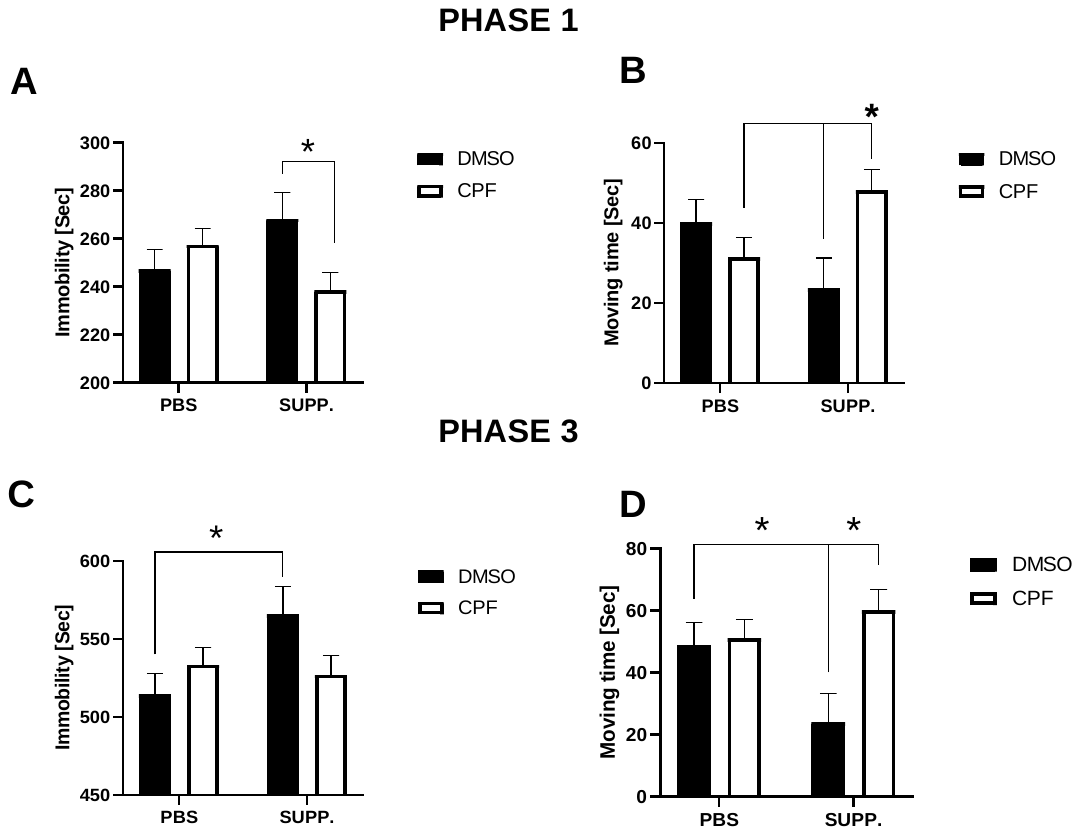

**Appendix 3B.** Locomotion in 3-Chambered Test. Means (±SEM) of immobility and movement in the habituation phase are presented (A and B), as well as the immobility and movement in the social novelty phase (C and D). *p<0.05

**Appendix 4**

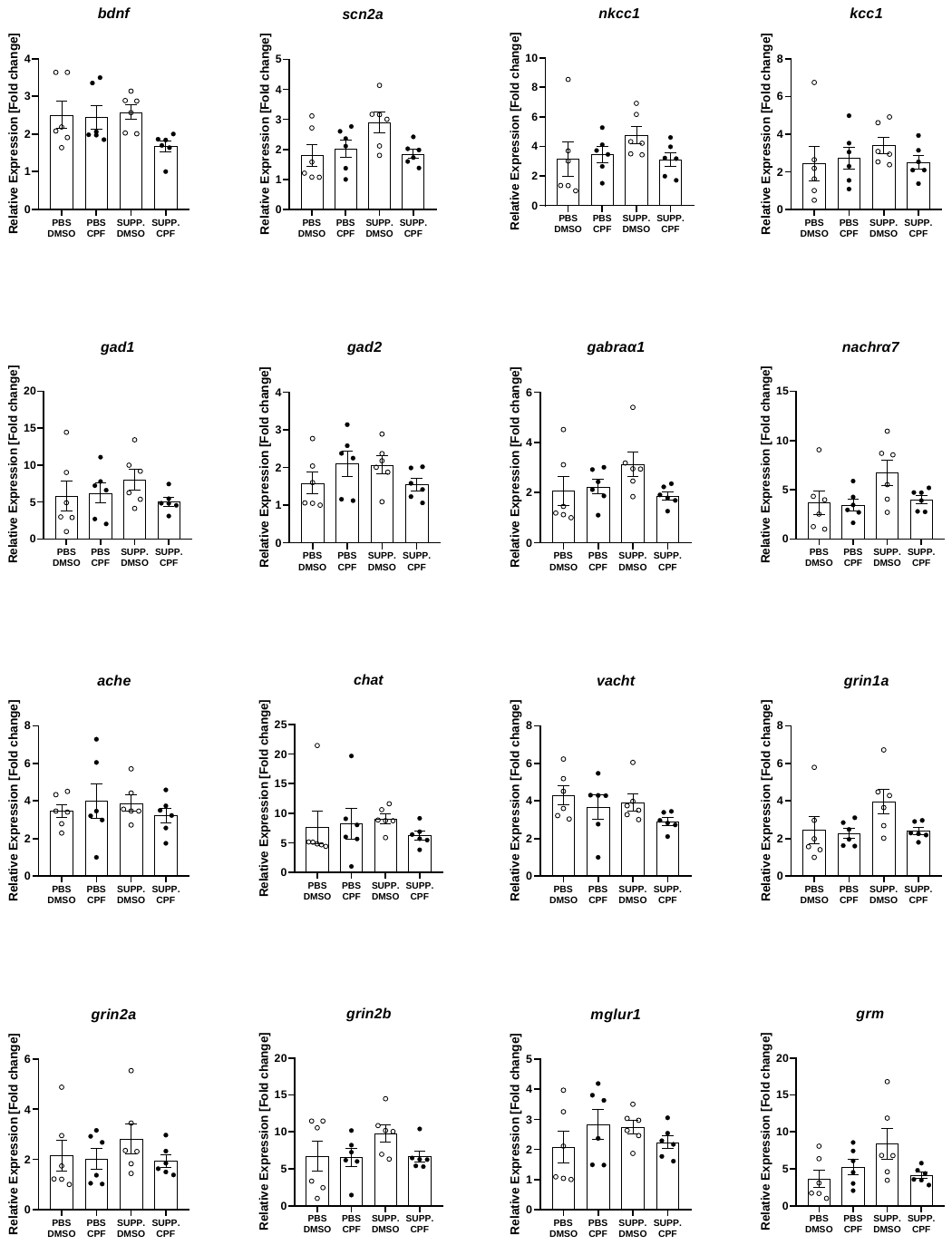

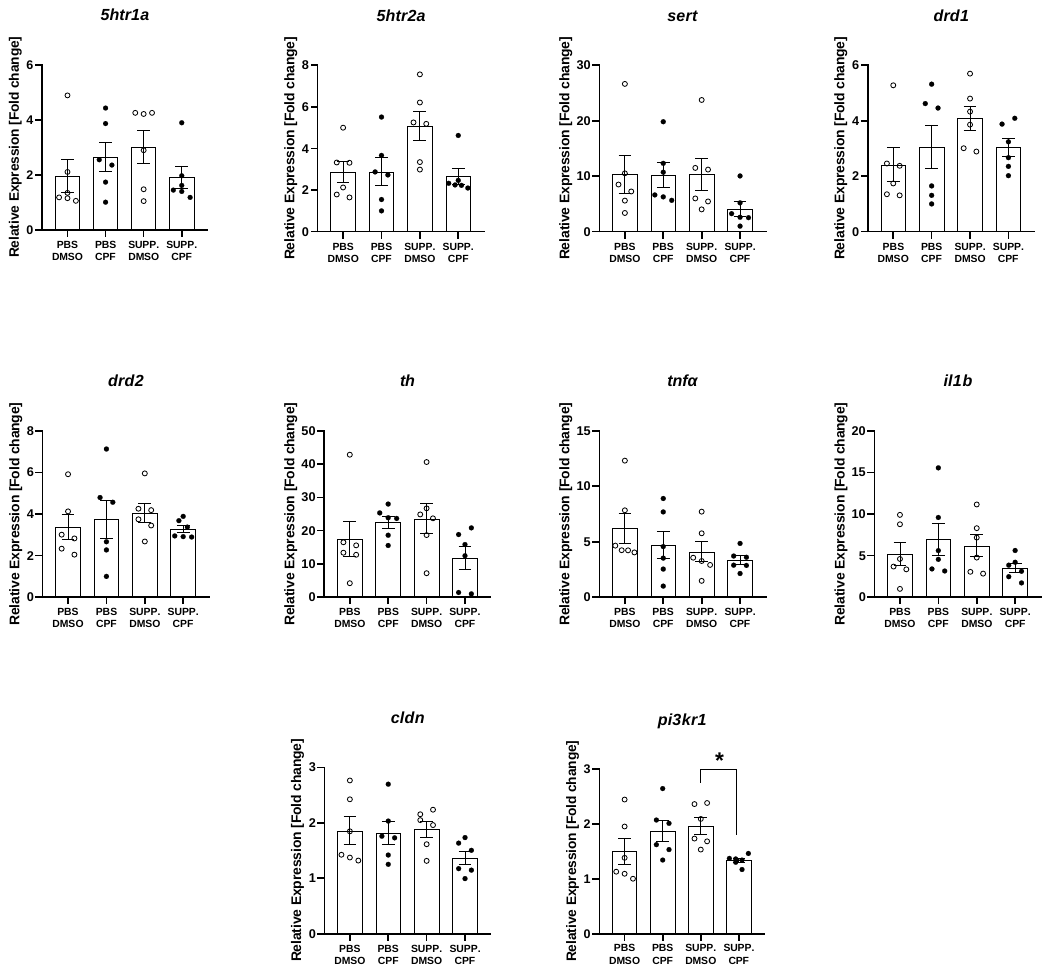

**Appendix 4.** Mean (±SEM) relative expression of all genes analysed in PND7 for which no statistically significant differences (p > 0.05) were found between groups. Also, *pi3kr1* expression has been represented (*p< 0.05).

**Appendix 5A**

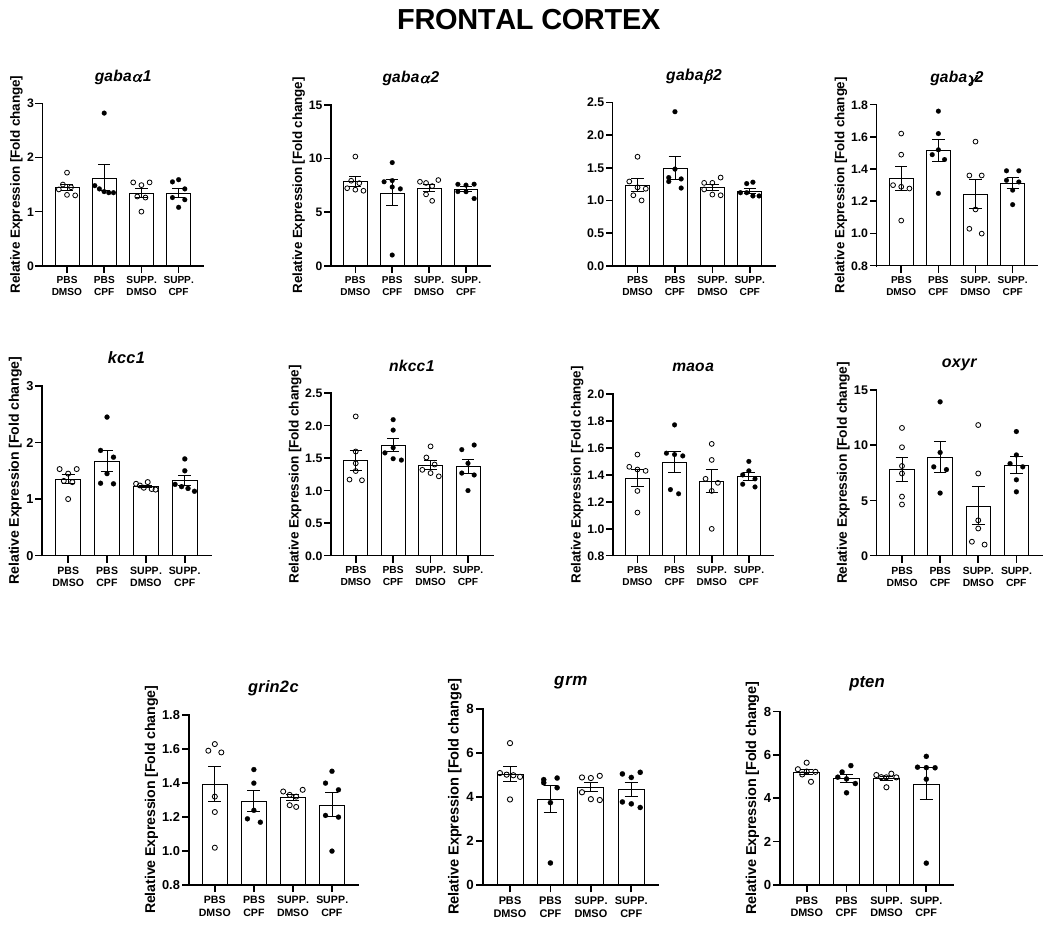

**Appendix 5.** Mean (±SEM) relative expression of all genes analysed in adolescence for which no statistically significant differences (p > 0.05) were found between groups (frontal cortex).

**Appendix 5B**

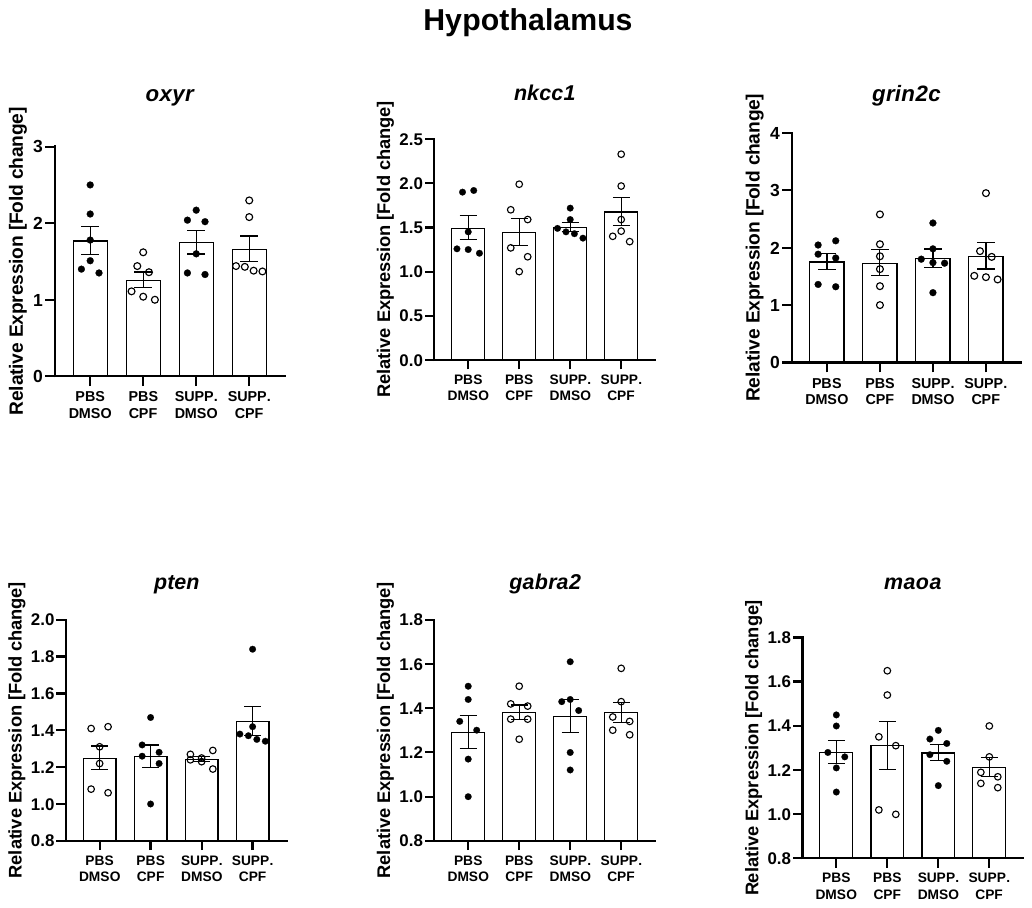

**Appendix 5B.** Mean (±SEM) relative expression of all genes analysed in adolescence for which no statistically significant differences (p > 0.05) were found between groups (hypothalamus).

**Appendix 6**

| Protein | Gene | Forward | Reverse | Source |
| --- | --- | --- | --- | --- |
| Glyceraldehyde-3-Phosphate Dehydrogenase  Interleukin 1 beta  Tumoral Necrosis Factor alpha  Vasopressin Receptor Subfamily 1a  Oxytocin Receptor  Monoamine Oxidase A  Acetylcholinesterase  Cholinergic Receptor Nicotinic subunit a7  Choline O-Acetyltransferase  Vesicular Acetylcholine Transporter  Glutamate Decarboxylase 2  Glutamate Decarboxylase 1  GABA Receptor Subunit alfa 1  GABA Receptor Subunit alfa 2  GABA Receptor Subunit beta 2  GABA receptor subunit gamma 2  Glutamate Metabotropic Receptor 2  Glutamate Receptor Subunit 2c  Glutamate Receptor Subunit 2b  Glutamate Receptor Subunit 2a  Glutamate Receptor Subunit 1a  Glutamate Metabotropic Receptor 1  Phosphoinositide-3-Kinase Regulatory Subunit 1  Phosphatase and tensin homolog  Forkhead Box P1  Dopamine Receptor D1  Dopamine Receptor D2  Tyrosine Hydroxylase  Serotonin receptor 2A  Serotonin receptor 1A  Serotonin Transporter  KCC2  KCC1  NKCC1  Sodium Voltage-Gated Channel Alpha Subunit 2  Claudin-5  Brain-Derived Neurotrophic Factor | *gapdh*  *il1b*  *tnfa*  *avpr1a*  *oxyr*  *maoa*  *ache*  *nachra7*  *chat*  *vacht (slc18a3)*  *gad2*  *gad1*  *gabra1*  *gabra2*  *gabrb2*  *gabrg2*  *grm (grm2)*  *grin2c*  *grin2b*  *grin2a*  *grin1a*  *mglur1(grm1)*  *pi3kr1*  *pten*  *foxp1*  *drd1*  *drd2*  *th*  *5htr2a*  *5htr1a*  *sert*  *slc12a5*  *slc12a4*  *slc12a2*  *scn2a*  *cldn5*  *bdnf* | ctgggtggctcaaggaata  cagctatggcaactgtccct  ggagggagaacagcaactcc  tacgtgacctggatgaccag  gagcgtttgggacgtcaatg  gtgtggaaccccttggcata  gtgagcctgaacctgaagcc  tatcaccaccatgaccctga  atggccattgacaaccatcttctg  gccacatcgttcactctcttg  ctgagaagccagcagagagc  gtgagtgccttcagggagag  gcccaataaactcctgcgtatc  ccaggatgacggaacattgc  gcacgttggagatcgaaagc  gaaaaaccctgcccctacca  ctatgccacccacagtgatg  ggcccagcttttgaccttagt  aagttcacctatgacctttacc  agttcacctatgacctctacc  atggcttctgcatagacc  agctgtgtctacccggactc  agccattgagaagaaaggactgg  acaaagacaaggccaaccga  ccctctgtcatcaccaccac  gcatggcttggattgctacg  cagtcgagctttcagagcca  ccttccagtacaagcacggt  aacggtccatccacagag  gatctcgctcacttggctca  ccgtcatctgcatccctacc  aggtggaagtcgtggagatg  catgatttcccgctctttg  catggtgtcaggatttgcac  tgcactggagactgctacat  cggaaagaccgatgtgggaa  ggtcacagcggcagataa | cacacgcatcacaaaaaggt  catctggacagcccaagtca  gccagtgtatgagagggacg  agcaacgccgtgattgtgat  acgagttcgtggaagaggtg  cccattcctgagcgtgtctt  tcctgcttgctatagtggtc  cagaaaccatgcacaccagt  aacaaggctcgctcccacagcttc  cggttcatcaagcaacacatc  agagtgggcctttctccttc  cgtcttgcggacatagttga  attcggctctcacagtcaacct  ggaaagtcctccaagtgcatt  cgtgactgcattgtcatcgc  tgcgaatgtgtatcctcccg  gcacagtgcgagcaaagtaatc  cctgtgaccaccgcaagag  catgaccacctcaccgat  gttgatagaccacttcacct  gttgtttacccgctcctg  ccgaaggtgtgtggatcagg  acgtgtacatcgaacacatcca  agcctctggatttgatggctc  ggtggtctaacttccgcgtt  ccagttgctgcctggactaa  ccaattctccgcctgttcact  tgggtagcatagaggccctt  aacaggaagaacacgatgc  aaagcgccgaaagtggagta  atgtccccacacgggatttc  cgagtgttggctggattctt  ccgtacacccgcatgttatt  gatattgtccttacatagag  ctcgcgtaagaaagtgctga  acccaacctaacttgcctcg  ccgaacatacgattgggtag | Own design  Own design  Own design  Own design  Own design  Own design  Jameson et al. (2007)  Chamoun et al. (2016)  Lips et al. (2007)  Lips et al. (2007)  Own design  Own design  Fujimura et al. (2005)  Fujimura et al. (2005)  Own design  Ruiz-Sobremazas et al. (2023)  Pershina et al. (2019)  Lau et al. (2013)  Lau et al. (2013)  Lau et al. (2013)  Lau et al. (2013)  Own design  Own design  Own design  Own design  Own design  Own design  Own design  Kindlundh-Högbergetal et al. (2006)  Own design  Own design  Jaenisch et al. (2010)  Own design  Cho et al. (2013)  Own design  Own design  Own design |

**Appendix 6.** Primers selected for the RTqPCR analysis.

**Table S1.** Metabolites identified in gut extracts from PND7 rats by ^1^H NMR

| Metabolite | δ_H_ (ppm), *J* (Hz) |
| --- | --- |
| 1. Bile acids | 0.71 (b.s.) |
| 2. Sterols | 0.74 (s), 0.79 (s), 0.81 (s) |
| 3. Fatty acids | 0.88 (t), 1.29 (m), 1.56 (m), 2.14 (m) |
| 4. Valine | 1.02 (d, *J* = 7.0 Hz), 1.07 (d, *J* = 7.0 Hz) |
| 5. Isoleucine | 0.96 (t, *J* = 7.2 Hz), 1.04 (d, *J* = 7.0 Hz) |
| 6. Leucine | 0.97 (m) |
| 7. 3-hydrobybutyrate | 1.20 (d, *J*= 6.3 Hz) |
| 8. Lactate | 1.34 (d, *J* = 6.9 Hz), 4.05 (q, *J* = 6.9 Hz) |
| 9. Alanine | 1.50 (d, *J* = 7.2 Hz) |
| 10. Arginine | 1.68 (m), 3.02 (t) |
| 11. Acetate | 1.91 (s) |
| 12. Proline | 1.96 (m), 2.04 (m), 2.34 (m) |
| 13. Glutamate | 2.06 (m), 2.39 (m) |
| 14. Glutamine | 2.14 (m), 2.46 (m) |
| 15. Succinate | 2.41 (s) |
| 16. Aspartate | 2.66 (dd, *J* = 17.5; 9.3 Hz), 2.80 (dd, *J* = 17.5; 3.6 Hz) |
| 17. Sarcosine | 2.71 (s) |
| 18. N,N-dimethylglycine | 2.93 (s) |
| 19. Creatine | 3.05 (s), 3.90 (s) |
| 20. Ethanolamine | 3.12 (t) |
| 21. Taurine | 3.21 (t, *J* = 6.3 Hz), 3.38 t, *J* = 6.3 Hz) |
| 22. Choline | 3.22 (s) |
| 23. Choline-derivative | 3.25 (s) |
| 24. Betaine | 3.27 (s) |
| 25. Glycine | 3.51 (s) |
| 26. Glycerol | 3.55 (dd, *J*= 6.2 Hz, 11.6 Hz), 3.62 (dd, *J*= 4.6, 11.6 Hz) |
| 27. Glucose | 4.62 (d, *J*= 7.5 Hz), 5.20 (d, *J*= 3.8 Hz) |
| 28. Sucrose | 4.19 (d, *J*= 8.9 Hz),5.43 (d, *J*= 3.6 Hz), |
| 29. Fumarate | 6.54 (s) |
| 30. Tyrosine | 6.85 (d, *J* = 8.5 Hz), 7.19 (d, *J* = 8.5 Hz) |
| 31. Phenylalanine | 7.40 (m), 7.34 (m) |
| 32. Uridine | 5.95 (d, J = 8.1 Hz), 5.96 (d, *J* = 4.6 Hz), 7.99 (d, *J* = 8.2 Hz) |
| 33. Xanthine | 7.99 (s), 5.97 (d), 5.96 (d) |
| 34. Formate | 8.50 (s) |
| 35. Nicotinate | 8.96 (s), 8.58 (m), 8.35 (m) |
| 36. Methylnicotinate | 9.52 (s), 9.32 (d, *J* = 6.6 Hz), 8.95 (d, *J* = 6.6 Hz), 8.30 (m) |

**Table S2.** Metabolites identified in fecal extracts from in PND37 rats by 1H NMR

| Metabolite | δ_H_ (ppm), *J* (Hz) |
| --- | --- |
| 1. Bile acids | 0.71 (b.s.) |
| 2. Sterols | 0.74 (s), 0.79 (s), 0.81 (s) |
| 3. n-butyrate | 0.90 (t), 1.56 (m), 2.17 (t) |
| 4. Proprionate | 1.06 (t), 2.20 (q) |
| 5. Valine | 1.02 (d, *J* = 7.0 Hz), 1.07 (d, *J* = 7.0 Hz) |
| 6. Isoleucine | 0.96 (t, *J* = 7.2 Hz), 1.04 (d, *J* = 7.0 Hz) |
| 7. Leucine | 0.98 (m) |
| 8. Alanine | 1.50 (d, *J* = 7.2 Hz) |
| 9. Arginine | 1.68 (m), 3.02 (t) |
| 10. Proline | 1.96 (m), 2.04 (m), 2.34 (m) |
| 11. Acetate | 1.91 (s) |
| 12. Glutamate | 2.06 (m), 2.39 (m) |
| 13. Glutamine | 2.14 (m), 2.48 (m) |
| 14. Succinate | 2.41 (s) |
| 15. Aspartate | 2.66 (dd, *J* = 17.5; 9.3 Hz), 2.80 (dd, *J* = 17.5; 3.6 Hz) |
| 16. Sarcosine | 2.71 (s) |
| 17. N,N-dimethylglycine | 2.93 (s) |
| 18. Choline | 3.22 (s) |
| 19. Choline-derivative | 3.25 (s) |
| 20. Betaine | 3.27 (s) |
| 21. Glycine | 3.51 (s) |
| 22. Glucose | 4.62 (d, *J*= 7.5 Hz), 5.20 (d, *J*= 3.4 Hz) |
| 23. Tyrosine | 6.85 (d, *J* = 8.5 Hz), 7.19 (d, *J* = 8.5 Hz) |
| 24. Phenylalanine | 7.40 (m), 7.34 (m) |
| 25. Formate | 8.48 (s) |
